# A single MHCII neoepitope mRNA vaccine elicits CD4 T- and B- cell responses promoting endogenous CD8 anti-tumor immunity

**DOI:** 10.1101/2025.06.09.657934

**Authors:** Aude-Hélène Capietto, Camille-Charlotte Balança, Romain Bouziat, Thomas D. Wu, Joshua Gober, Catherine Carbone, Ariane Nissenbaum, Reyhane Hoshyar, Marshleen Yadav, Alan Gutierrez, Yajun Chestnut, Vincent Javinal, Hari Menon, Spyros Darmanis, Marco de Simone, Yonglian Sun, Dhaya Seshasayee, Maciej Paluch, Beyza Bulutoglu, Tamaki Jones, Yingyun Wang, Lisa Liao, Ellen Duong, Aditya Anand, Anthony Antonelli, Yoko Oei, Jeanne Cheung, Shiuh-Ming Luoh, James Ziai, Jessica Preston, Jeffrey Hung, Emily Freund, Sara Wichner, Jaehak Oh, Siri Tahtinen, Ugur Sahin, Adel ElSohly, Jill Schartner, Lélia Delamarre, Ira Mellman, Jonathan L. Linehan

## Abstract

Recent progress in therapeutic cancer vaccines has shown promising clinical activity, especially when targeting MHC class I (MHCI) neoantigen-specific CD8^+^ T cell responses in post-surgical patients. To explore the role of CD4^+^ T cells in vaccine-dependent tumor rejection, we constructed an mRNA lipoplex vaccine encoding a single MHCII-restricted neoantigen. The vaccine elicited Tfh and Th1 cell responses while decreasing Tregs, leading to rejection of established tumors in mice. IL-21 and IFN-γ, crucial for Tfh and Th1 function respectively, contributed to anti-tumor activity. B cells and neoantigen-specific antibodies were also shown to participate in vaccine efficacy. Additionally, conventional type 1 dendritic cells (cDC1s) were essential for eliciting vaccine-induced CD4^+^ T cells, and both cDC1s and CD4^+^ T cells were required to enhance endogenous CD8^+^ responses, which were crucial for tumor control. Our results suggest that immunizing against MHCII neoantigens alone is sufficient to orchestrate a potent and cooperative immune response against cancer.

## Introduction

Among the emerging strategies in cancer immunotherapy, individualized mRNA-based vaccines have garnered significant attention for their ability to stimulate a robust tumor-specific immune response by targeting patient-specific immunogenic tumor mutations, termed neoantigens^1–5^.

Although the focus of cancer vaccines has mainly been on targeting CD8^+^ T cells, there is substantial evidence to support the importance of eliciting CD4^+^ T cell responses. Vaccination using only MHCII-restricted CD4 neoantigens, or adoptive transfer of neoantigen-specific CD4^+^ T cells, have been shown to confer efficient anti-tumor responses in mice and humans^6–8^. The underlying mechanisms are not yet clear, however, and may involve the direct killing of MHCII-positive tumor cells by cytotoxic CD4^+^ T cells, cytokine-induced (e.g. IFN-γ) tumor cell death, or “help” provided by CD4^+^ T cells in the priming and maintenance of CD8^+^ T cells, B cells, and innate immune cells^9, 10^.

CD8^+^ T cells serve as the primary cytotoxic effector cells responsible for the destruction of tumor cells. Effective and long-lasting cytotoxic activity, especially in the context of immunotherapies, relies on the cooperation between CD4^+^ and CD8^+^ T cells in both lymphoid tissues and tumors^10^. In the priming phase, CD4^+^ T cells help conventional dendritic cells (cDCs) to activate CD8^+^ T cells, partially through CD40/CD40L signaling, and to facilitate the clonal expansion and differentiation of CD8^+^ T cells into effector and memory, and cytotoxic T lymphocytes (CTLs)^9, 11, 12^. Considerable evidence supports a major role for the cDC1 population in the cross presentation of tumor antigens on MHCI molecules to CD8^+^ T cells, with cDC2s being more associated with priming CD4^+^ T cells^13, 14^. However, in the context of viral and tumor antigens, cDC1s have a greater capacity than cDC2s for processing cell-associated MHCII-restricted antigens^15, 16^. Furthermore, work by Ferris et al.^17^ demonstrated that cDC1s can prime tumor-specific CD4^+^ T cells and are subsequently licensed by CD4^+^ T cells to optimize anti-tumor CD8^+^ T cell responses. Conversely, recent evidence has suggested that cDC2s can also efficiently cross-present MHCI-restricted antigens to CD8^+^ T cells, at least when antigens were captured by Fcγ receptor-mediated endocytosis^18^.

In addition to lymphoid tissues, CD4^+^ T cells provide help to CD8^+^ T cells in the tumor microenvironment (TME) for efficient spontaneous and immune checkpoint blockade (ICB)-mediated anti-tumor responses^19^. Particularly, recent studies have suggested the importance of CD4^+^-cDC interactions within the TME to maintain cytotoxic CD8^+^ T cell responses in mouse models and human hepatocellular carcinoma^20, 21^. Although these studies demonstrate the formation of “triads” comprising DCs, CD4^+^, and CD8^+^ T cells as being associated with CD8^+^ T cell responses, the exact mechanisms as well as the nature of cDC subsets and CD4^+^ T cells involved remain unclear, especially in the context of cancer vaccines.

Many studies of neoantigen-targeted therapies report a predominant T helper 1 (Th1) cell phenotype among vaccine-induced CD4^+^ T cells^19, 22, 23^. The specific roles of individual CD4^+^ subsets in anti-tumor immunity remain to be elucidated, however^24^. Emerging evidence that tumor-specific antibodies in cancer patients and tumor-infiltrating B cells may be associated with favorable outcomes suggests that T follicular helper (Tfh) cells, given their role in enhancing B cell activity and generating effective antibody responses^25^, may also play significant roles in cancer vaccines^26^. Here, we demonstrate that a tumor-specific MHCII-restricted neoantigen vaccine expands CD4^+^ Tfh and Th1 cells, improving B cell responses and expanding plasma cell clones that produce neoantigen-specific antibodies; B cells were also found to be required for vaccine-induced anti-tumor activity. Importantly, cDC1s prime CD4^+^ T cells in the context of a neoantigen encoded mRNA vaccine and facilitate CD4^+^ T cell help to endogenous CD8^+^ TILs, responsible for tumor rejection. Our findings underscore the critical function of this DC subset in vaccine-induced immune responses. As the vaccine did not contain MHCI epitopes, our data demonstrate that eliciting a CD4^+^ response can be sufficient to induce endogenous CD8^+^ responses to tumor antigens presented by cDC1s. Altogether, the RNA-LPX vaccine encoding for a single MHCII-restricted neoantigen, successfully initiates a strong and coordinated immune response, leading to tumor elimination.

## Results

### MHCII-restricted neoantigen vaccination induces E0771 tumor rejection, independent of MHCII expression on tumor cells

To investigate the role and mechanisms of tumor-specific CD4^+^ T cells in vaccine therapy, we utilized the E0771 mammary carcinoma model, which features a previously identified MHCII-restricted neoantigen, designated envRV^27^. This neoantigen is derived from the envelope protein of Murine Leukemia virus (MuLV) and contains two point mutations. Vaccination with an envRV peptide vaccine has been shown to delay E0771 tumor growth^27^.

To monitor and characterize envRV-specific CD4^+^ T cells, we developed a pMHCII tetramer^28^ (Figure S1A-C). To provide for a control MHCII peptide, we used the well-characterized 2W1S:I-Ab epitope, which is known to elicit a robust CD4^+^ T cell response but is not expressed by E0771 tumor cells^29–31^ (Figure S1C). After confirming that RNA-LPX vaccines encoding either envRV and 2W1S exclusively induced MHCII-restricted CD4^+^ T cell responses (Figure 1A), we evaluated their anti-tumor activity. We vaccinated mice either on the same day as tumor inoculation or 7 days later when tumors were established. The neoantigen-specific envRV vaccine induced tumor rejection in both scenarios, whereas tumors grew continuously in unvaccinated mice or mice receiving the control 2W1S vaccine (Figure S1D and Figure 1B). Notably, envRV RNA-LPX vaccination resulted in an 11-fold increase in envRV-tetramer positive cells in both spleen and tumor compared to the non-vaccinated animals (Figure 1C). Although comparable or even higher frequencies of 2W1S-specific CD4^+^ T cells were observed in spleens and tumors of 2W1S RNA-LPX vaccinated animals (Figure 1C), no anti-tumor effect was noted (Figure 1B), consistent with the lack of 2W1S antigen expression by E0771 tumor cells.

**Figure 1:**
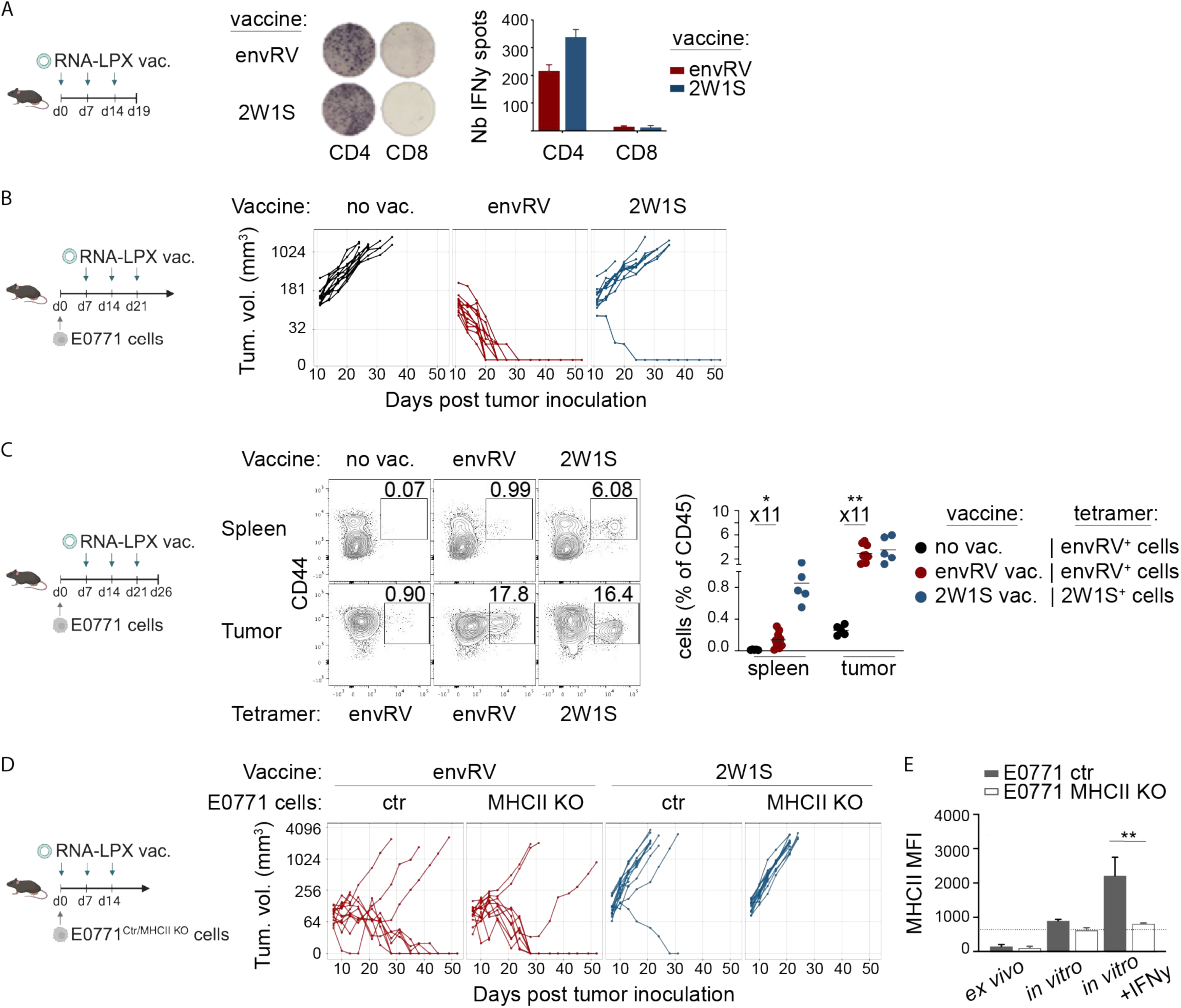
EnvRV RNA-LPX elicits a specific CD4^+^ T cell response and induces E0771 tumor rejection independent of tumor cell MHCII expression. A: Mouse IFN𝛾 ELISpot of splenocytes depleted of CD8^+^ T cells (CD4 response) or CD4^+^ T cells (CD8 response) from vaccinated mice with envRV- or 2W1S-RNA-LPX. 5x10^5^ CD4 or CD8 depleted-splenocytes were restimulated overnight with either envRV or 2W1S peptides. Representative ELISpot images and quantification of the number of IFN𝛾 spots for each condition are shown. Data are represented as mean +/- SD. B: E0771 tumor-bearing mice were vaccinated either with envRV (neoantigen) or 2W1S (irrelevant tumor antigen) RNA-LPX starting 7 days post tumor inoculation. A total of 3 vaccines, 7 days apart, were administered. Non-vaccinated animals (no vac.) were used as control. Volume of tumors were measured for 52 days post tumor cell inoculation. C: EnvRV- and 2W1S-specific tetramer staining in spleen and tumor of envRV-(neoantigen) and 2W1S- (irrelevant tumor antigen) vaccinated animals. Non-vaccinated animals were used as controls and stained with envRV tetramers. Representative FACS plots and quantification of the mean percentage of tetramer^+^ cells for all conditions are shown. Fold changes (x11) between the percentages of envRV^+^ CD4^+^ T cells in the no vaccine and envRV vaccinated groups for spleen and tumor samples are indicated on the graph. EnvRV^+^ cells in the no vaccine and envRV vaccinated group were compared using a multiple unpaired t test and showed statistical significance in both tissues (spleen, p = 0.02; tumor, p = 0.001). See also Figure S1. D: Mice were inoculated with either E0771 control (ctr) or E0771 MHCII KO cells and vaccinated with envRV (neoantigen) or 2W1S (irrelevant tumor antigen) RNA-LPX starting on the same day as tumor inoculation. A total of 3 vaccines, 7 days apart, were administered. Volume of tumor was measured for 52 days post tumor cell inoculation. E: E0771 (ctr, grey bar) or E0771 MHCII KO (white bar) cells were stained for MHCII surface expression *ex vivo* or after 72 hours of culture with or without IFN𝛾 (*in vitro*). Data are represented as mean of MHCII MFI +/- SEM.

Since neoantigen-specific CD4^+^ T cells have been shown to directly kill tumor cells in an MHCII-restricted manner^32, 33^, we next engineered E0771 cells lacking MHC-II expression to test whether the envRV RNA-LPX vaccine would still protect against tumor growth. In contrast to the control vaccine, E0771 tumors were similarly rejected following envRV vaccination independent of MHCII tumor cell expression (Figure 1D-E), indicating that the vaccine-elicited tumor-specific CD4^+^ T cells do not directly kill E0771 tumor cells through cognate interactions.

### cDC1s prime CD4^+^ T cells following MHCII-restricted antigen RNA-LPX vaccination

Intravenous administration of RNA-LPX vaccine mainly targets the spleen and generates T cell responses that depend on antigen presentation by cDCs^34^. However, it remains unknown which cDC subset(s) is responsible for T cell priming in the context of MHCII-restricted antigen RNA-LPX.

Using *Batf3^-/-^* cDC1-deficient mice, we first assessed whether cDC1s play a role in controlling tumor growth following MHCII-neoantigen vaccination. In comparison to WT mice, the envRV RNA-LPX vaccine failed to reject E0771 tumors in *Batf3^-/-^* mice, demonstrating the critical role of cDC1s in MHCII neoantigen vaccine efficacy (Figure 2A). Surprisingly, both envRV- and 2W1S- specific CD4^+^ T cells were significantly reduced in spleen and tumor from vaccinated *Batf3^-/-^* mice (Figure 2B) and the presence of cDC2s did not compensate for the generation of vaccine-induced CD4^+^ T cells. This result was unexpected given that: i) both cDC1s and cDC2s internalized the RNA-LPX vaccine, efficiently expressed mRNA-LPX encoded antigens, and presented antigens on MHCII (Fig S2A-B), and ii) cDC2s are generally considered to preferentially prime CD4^+^ T cells^35, 36^. This result demonstrates that cDC1s are the primary drivers of CD4^+^ T cell priming following RNA-LPX vaccination.

**Figure 2:**
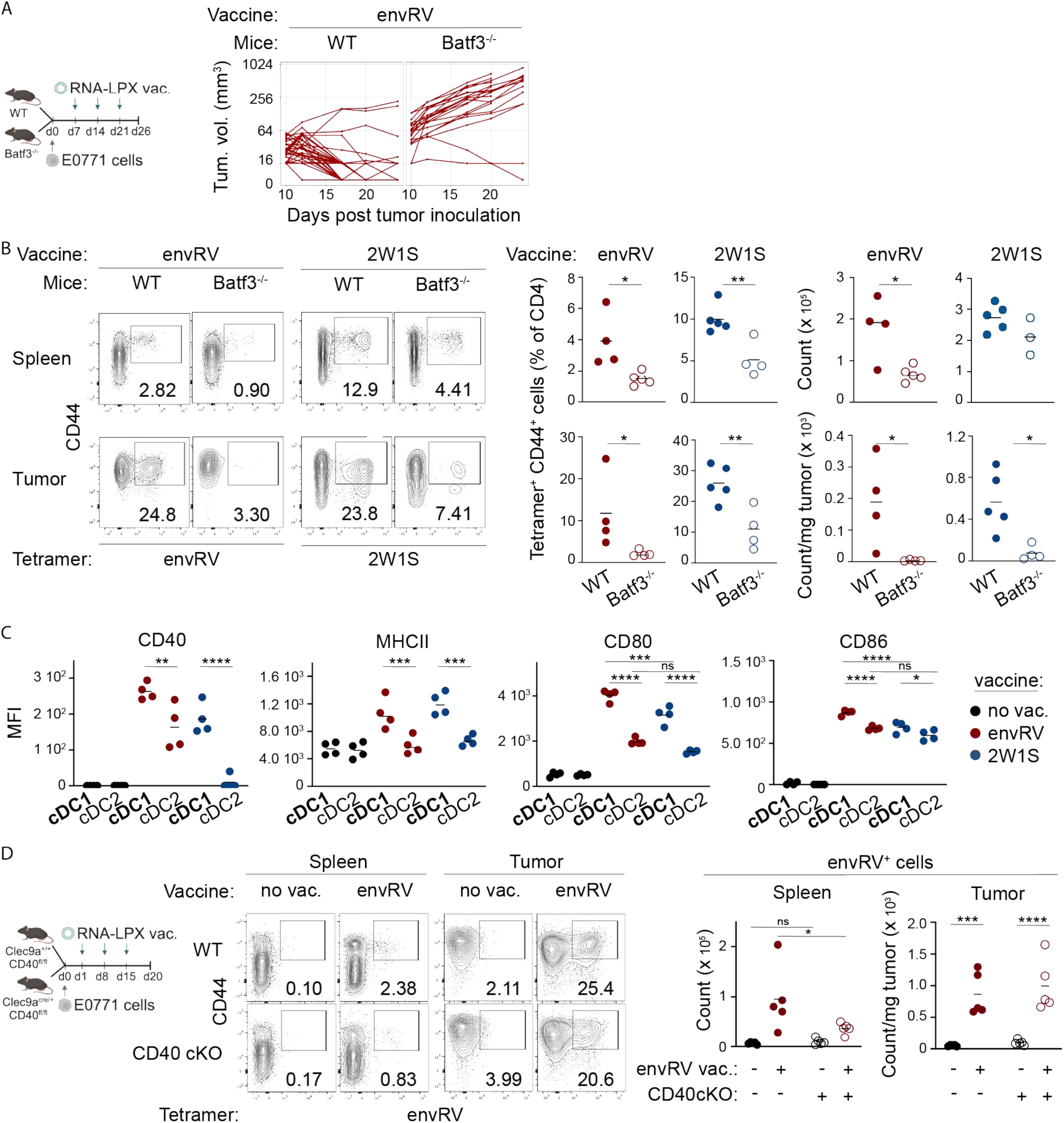
cDC1s are largely contributing to vaccine-specific CD4^+^ T cell priming and vaccine-induced anti-tumor efficacy. A: Study design and tumor growth curves of WT and Batf3^-/-^ mice vaccinated with envRV RNA-LPX. B: Flow cytometry analysis of tetramer^+^CD44^+^ CD4^+^ T cells in spleen (top) and tumor (bottom) from WT and Batf3^-/-^ mice 26 days post tumor inoculation. Representative FACS plots are shown (left). Mean of proportion and number of envRV^+^ and 2W1S^+^ cells among CD4^+^ T cells are shown for one representative experiment out of 1 to 3 independent studies (envRV, spleen n=3; envRV, tumor n=2; 2W1S, spleen and tumor n=1). Data are shown from two different experiments for envRV and 2W1S vaccines. Normality and Mann-Whitney or unpaired t tests were used. *p<0.05, **p<0.01. C: MFI of CD40, MHCII, CD80 and CD86 surface expression on cDC1s and cDC2s from spleen of WT tumor-bearing mice 24 hours post last vaccination. WT mice were vaccinated starting 7 days post E0771 tumor cell inoculation for a total of 3 times, 7 days apart. Data shown as mean from one representative experiment out of 4 independent studies. ns: non significant, *p<0.05, **p<0.01, ***p<0.001, ****p<0.0001. D: Clec9acre^+^/^+^ CD40^fl/fl^ (CD40 cKO) and Clec9a^+^/^+^ CD40^fl/fl^ littermate (control) mice were vaccinated or not with the envRV RNA-LPX vaccine 7 days apart starting one day post E0771 tumor inoculation. Spleens and tumors were harvested 5 days post last vaccine for flow cytometry analysis. Representative FACS plots showing tetramer staining of envRV^+^ CD44^+^ CD4^+^ T cells in spleen and tumor. Data shown as mean of the count (spleen) or count/mg of tumor. ns: non significant, *p<0.05, ***p<0.001, ****p<0.0001. n=1 experiment.

Next, we analyzed the maturation status of splenic cDCs from vaccinated tumor-bearing mice. Interestingly, cDC1s showed higher expression of the maturation and activation molecules MHCII, CD80, CD86 and CD40 than cDC2s (Figure 2C), suggesting that RNA-LPX vaccines differentially activate the two cDC subsets. As CD40 signaling in cDC1s contributes to CD4^+^ T cell proliferation in mice immunized with cell-associated antigen^17^, we assessed envRV-specific CD4^+^ T cell generation in *Clec9a*^cre/+^*CD40*^fl/fl^ (CD40 cKO) tumor-bearing mice lacking CD40 expression in cDCs. EnvRV-specific CD4^+^ T cell number in spleen was significantly reduced in response to vaccination in CD40 cKO mice compared to littermate controls (Figure 2D). Nevertheless, the number of tumor-infiltrating envRV^+^ T cells was unchanged between the two groups (Figure 2D). Thus, CD40 signaling in cDCs is only partially involved in generating envRV-specific CD4^+^ T cells.

### EnvRV RNA-LPX vaccine promotes CD4^+^ Tfh and Th1 cell expansion to control tumor growth

We next examined how envRV RNA-LPX vaccination affected the phenotype and distribution of CD4^+^ T cells. We performed single-cell RNA sequencing (scRNAseq) and TCR sequencing (scTCRseq) on CD4^+^ T cells isolated from spleens and tumors of vaccinated animals (Figure S3A and Methods).

Gene expression analysis identified 14 clusters with distinct CD4^+^ T cell phenotypes (Figure 3A, B). We saw classifications of canonical helper T cell phenotypes including Tfh, Th1, regulatory T cell (Treg), and naive as well as non-canonical CD4^+^ T cell phenotypes (see details in Methods). We then tabulated cluster composition by organ and treatment. As expected, naive-like clusters were well established in spleen but limited in tumor. Conversely, the Treg cluster was present predominantly in tumors, but the size of this cluster was not altered by vaccination with either antigen (Figure 3C). The Th1 clusters were found in both organs and groups but to a greater extent in control samples than after envRV vaccination. On the other hand, while the Tfh cluster was limited and mostly restricted to spleen in the control group, the envRV vaccine caused a much greater accumulation of Tfh in both spleens and tumors (Figure 3C).

**Figure 3:**
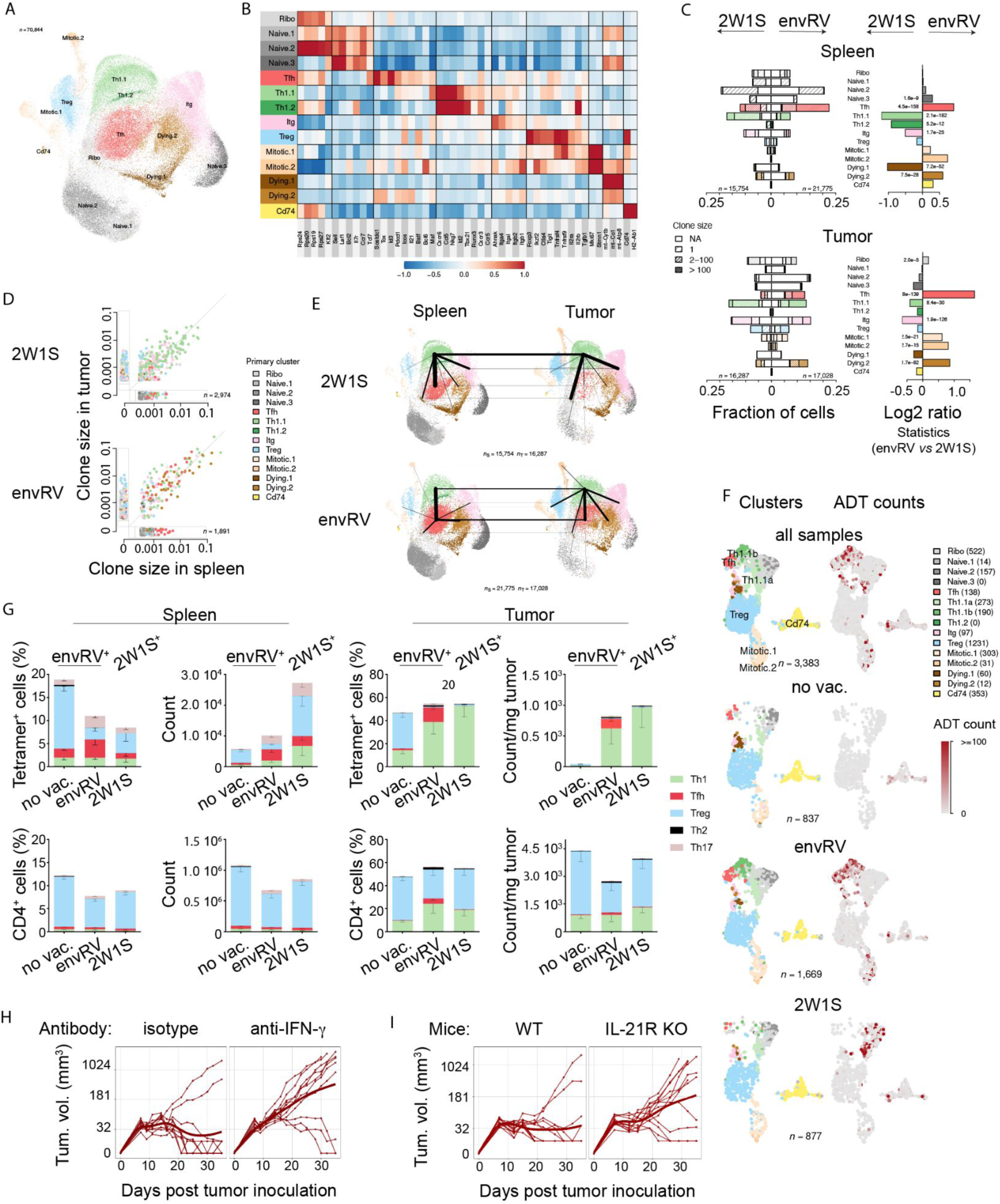
EnvRV RNA-LPX vaccine promotes expansion of CD4^+^ Tfh and Th1 cells A: UMAP of CD4^+^ T cells colored by phenotype (combination of spleens and tumors for the envRV- (n=3 samples each) and the 2W1S-vaccine groups (n=2 samples each); for a total of n=10 samples). Study design shown in Figure S3A. B: Heatmap showing relative average expression of selected marker genes associated with CD4^+^ T cell phenotype, function or differentiation states in each cluster identified in the UMAP in A. mt: mitochondrial. C: Stacked bar graphs of CD4^+^ T cell cluster composition as defined in UMAP for spleen (upper panel) and tumor (lower panel) tissues from control (2W1S, left side) and envRV (right side) RNA-LPX vaccinated mice. In each stacked bar, open bar denotes cells for which no TCR expression has been identified, light stripes for singletons, medium stripes for clones between 2-100 cells, and high density stripes for clones greater than 100 cells. Stripe color represents the primary cell phenotype of each clone as shown in the UMAP in A. Total number of cells are shown for each condition: 2W1S/spleen, n=15754; envRV/spleen, n= 21775; 2W1S/tumor, n=16287; envRV/tumor, n=17028. Log2 ratio between 2W1S and envRV (envRV *vs* 2W1S) with statistically significant values are noted for both spleen and tumor tissues. D: Scatterplots showing the size of each individual clonotype in tumor and spleen for each vaccination treatment (2W1S, upper panel and envRV, lower panel). Color of circles denotes the primary cluster designation of each clone. Number of cells for each condition: n=2974 cells for 2W1S and n=1891 cells for envRV. See also Figure S3B. E: Cluster co-occurrence analysis in spleens (left panel) and tumors (right panel) from 2W1S (control, upper panel) and envRV (lower panel) vaccinated animals. Lines within UMAPs denote co-occurrences between different clusters within the indicated tissue. Lines between UMAPs denote co-occurrences between the same cluster in different tissues. Thickness of line denotes relative strength of co-occurrence, with thickest lines indicating strongest co-occurrence. Numbers of pairwise co-occurences per group are: 2W1S/spleen, n=15754; 2W1S/tumor, n=16287; envRV/spleen, n=21775; envRV/tumor, n=17028. Clusters are colored as in the UMAP in A. F: UMAP showing the transfer of CD4^+^ T cell analysis from the CD45^+^ scRNAseq dataset (detailed in Figure S3D-E) to the primary CD4^+^ scRNAseq described in Figure 3A-E. CD4^+^ T cell clusters colored by phenotype as in A for all samples (n=3383 cells), no vaccine (no vac., n=837 cells), envRV (n=1669 cells) and 2W1S (n=877 cells) RNA-LPX vaccines (left panels). Teramer^+^ CD4^+^ T cells are shown as ADT counts (depicted as red dots) in UMAP for each group; envRV^+^ cells for no vaccine and envRV RNA-LPX groups, 2W1S^+^ cells for the 2W1S RNA-LPX group (right panels). See also Figure S3F. G: Phenotypic analysis (study design in Figure S3A) of CD4 tetramer-positive cells (upper panel) and tetramer-negative cells (lower panel) in spleen and tumor of envRV or 2W1S vaccinated animals. No vaccination was used as control. EnvRV-specific cells were analyzed in non-vaccinated and envRV-vaccinated samples. 2W1S-specific cells were analyzed in 2W1S vaccinated samples. Cumulative bar graphs representing the frequency or count of the 5 phenotypes within the tetramer^+^ or CD4^+^ T cell populations. Data are shown as mean +/- SEM. H: E0771 tumor-bearing mice were vaccinated with envRV (neoantigen) RNA-LPX and received isotype control or anti-IFN-γ antibodies. Tumor volumes were measured for 35 days post tumor cell inoculation. See study design in Figure S3H. I: E0771 tumor-bearing WT or IL-21R KO mice were vaccinated with envRV (neoantigen) RNA-LPX. Tumor volumes were measured for 35 days post tumor cell inoculation. See study design in Figure S3I.

We then used the clonotypes from scTCRseq to measure clonal expansion in the various clusters. Most of the clones were found in both tumor and spleen (Figure 3D and S3B). The envRV group resulted in greater clonal expansion of both Tfh and Th1 clusters, while the expanded clones from the control group were mainly Th1 (Figure 3D). Many of the largest clones found in tumors of envRV-treated mice had similarly large matching ones in spleens (Figure S3B).

The mixed composition of phenotypes within clones provided us with an opportunity to study possible lineage relationships, since cells with the same TCR clonotype must share a common ancestor. By tracking phenotypic co-occurrences within clones, we can determine the most common differentiation links and infer primary tissue hubs and migration patterns using a minimum-spanning tree. For the 2W1S control group, we observed lineage relationship between the Th1 and Tfh phenotypes, which was more common in spleen; we also noted the predominant migration of the Th1 phenotype between spleen and tumor (Figure 3E, top panels). For the envRV-group, the primary hub of differentiation in tumor was also the Th1 phenotype, but more so in spleen; we also observed greater co-occurrence of Tfh cells within clusters, as well as accrued migration of the Tfh phenotype between spleen and tumor (Figure 3E, lower panels).

To validate that expanded CD4^+^ T cell clonotypes were vaccine-induced, we performed an additional experiment using barcoded-tetramers on intratumoral CD45^+^ cells and ADTseq, in addition to scRNA/TCRseq (Figure S3C). CD4^+^ T cells from this second dataset (Figure S3D-E) were projected onto the initial NGS analysis of total CD4^+^ T cells (Figure 3A-E). The projected cell types included all phenotypes but naive.3 and Th1.2 clusters. However, the projected cells in the Th1.1 cluster were divided into Th1.1a and b subclusters with the Th1.1b cluster sharing closer expression similarities to Tfh cells. Interestingly, only the envRV group contained substantial numbers of Tfh cells and Th1.1b cells while control groups mostly showed Th1.1a cells (Figure 3F, left panels). As confirmed with ADT sequencing, these Tfh and Th1.1b clusters specifically observed in the envRV group were envRV^+^ cells (Figure 3F, right panels and Figure S3F).

Next, we confirmed the phenotypes of envRV-specific cells by flow cytometry in both naive and tumor-bearing animals. We employed canonical transcription factors (TF) to identify each CD4^+^ T cell phenotype (Figure S3A).

Similar to the scRNAseq analysis, the frequency of GATA-3^+^ (Th2) and ROR𝛾t^+^ (Th17) cells was almost undetectable, regardless of vaccination status or organ type (Figure 3G). However, many CD4^+^ T cells were not accounted for with our gating strategy. This result may be explained by the plasticity of transient phenotypes (Figure 3E), possibly represented by the additional clusters observed in the scRNAseq analysis (naive, Itg, ribo, mitotic, CD74; Figure 3A-B). In non-vaccinated control mice, the few envRV-specific cells were mostly FoxP3^+^ Tregs in both spleens and tumors (Figure 3G, top panels). Interestingly, the frequency of envRV-specific Treg cells was significantly reduced post envRV vaccination in spleens and nearly abrogated in tumors, and replaced by Tfh and Th1 phenotypes. 2W1S-specific Tregs were also absent in tumors of vaccinated animals and tetramer-negative CD4^+^ TILs predominantly exhibited a Treg phenotype in all 3 groups (Figure 3G, lower panels). This result suggests that MHCII antigen-encoded RNA-LPX vaccines diminish the generation of intratumoral vaccine-induced Treg cells, independent of tumor-specificity. Importantly, while tumor-bearing mice vaccinated with envRV RNA-LPX developed both Th1 and Tfh responses, non-tumor-bearing mice only developed Th1 responses after vaccination (Figure S3G). This result suggests that the presence of the antigen in the tumor is driving Tfh expansion.

To determine the contribution of Th1 and Tfh subsets to envRV-vaccine induced efficacy, we examined the impact of blocking IFN-γ and IL-21 signalling pathways that are crucial to Th1 and Tfh cell function, respectively. Blocking IFN-γ, primarily produced by Th1 cells, abrogated envRV vaccine-induced efficacy (Figure 3H and S3H). Similarly, genetic deficiency of the IL-21 receptor (IL-21R) also severely blunted vaccine efficacy (Figure 3I and S3I). These results suggest that vaccine-elicited Th1 and Tfh responses were necessary for tumor rejection.

### Anti-tumor efficacy of envRV RNA-LPX vaccine is dependent on B cells and envRV-specific antibody

Since cognate interactions between Tfh cells and B cells are crucial for their mutual maturation and function, we next evaluated the importance of B cells in vaccine-induced tumor rejection using an anti-CD20 B cell-depleting antibody. Notably, B cell depletion clearly impaired tumor rejection following envRV RNA-LPX vaccination (Figure 4A). Further investigating the B cell compartment (Figure S4A), we found that GC, memory, and plasma B cell types were significantly increased in spleens of envRV vaccinated animals compared to unvaccinated controls, but not in tumors (Figure 4B).

**Figure 4:**
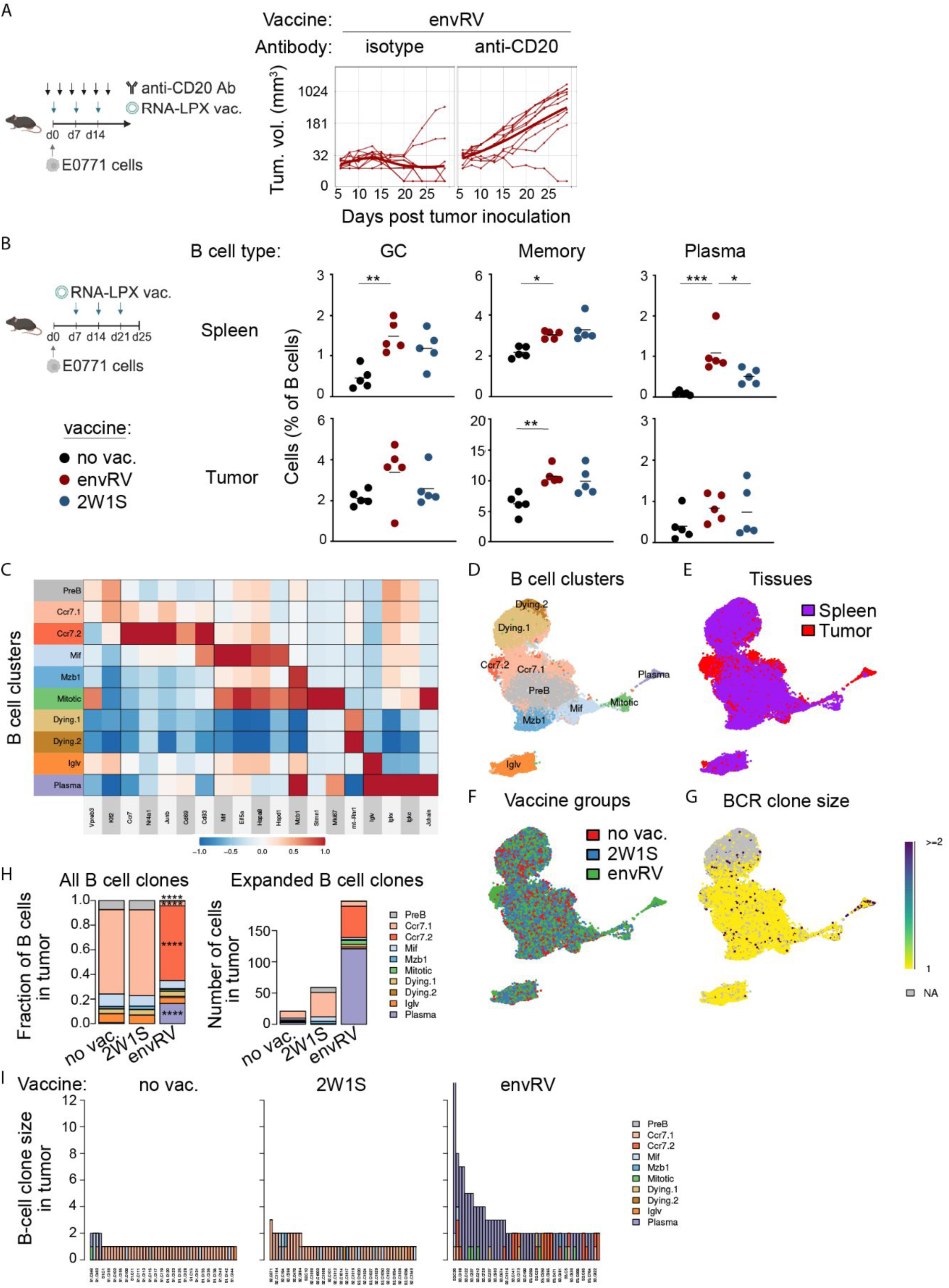
EnvRV RNA-LPX-induced antitumor efficacy is dependent on B cells. A: E0771 tumor-bearing mice were vaccinated with envRV (neoantigen) RNA-LPX starting at day 0 for a total of 3 vaccines given 7 days apart. Isotype control or anti-CD20 antibodies were administered at 10 mg/kg IV then 5 mg/kg IP biWx3 for a total of 6 injections. Volume of tumor was measured for 30 days post tumor cell inoculation. B: B cell phenotype analyzed by flow cytometry of spleens and tumors from mice bearing E0771 tumors and vaccinated with envRV or 2W1S RNA-LPX. Mean percentage of GC-, Memory-, and plasma-B cells are shown. *p<0.05; **p<0.01; ***p<0.001. See also Figure S4A. C: Heatmap showing relative average expression of selected marker genes associated with B cell phenotype, function or differentiation states in each cluster identified in the UMAP in D. B-cell subsets were identified in the CD45^+^ scRNA/BCRseq dataset. See also Figure S3D-E. D-G: UMAPs of B cells from spleens and tumors from the CD45^+^ scRNA/BCRseq dataset. (D) colored by clusters, (E) colored by tissue (spleen in purple and tumor in red), (F) colored by vaccine group (no vaccine in red (no vac.), 2W1S in blue, and envRV in green), (G) colored by clone size. H: Stacked bar graphs of B cell cluster composition of all cells (left) and expanded clones (right) in tumors from non-vaccinated (no vac.), control (2W1S) or envRV vaccinated mice. Significant statistical values between the envRV group and the two other groups are shown (****p<0.0001; pre-B, Ccr7.1, Ccr7.2, Mif and plasma clusters). I: Bar graph representing the size and phenotype of intratumoral B cell clonotypes from non-vaccinated (no vac.), control (2W1S) or envRV RNA-LPX vaccinated mice.

To better understand the phenotypic changes occurring in B cells following vaccination, we performed scRNAseq and scBCRseq on B220^+^ cells isolated from spleens and B cells computationally extracted from intratumoral CD45^+^ cells from the tumors of non-vaccinated or envRV/2W1S vaccinated animals (Figure S3C-D). Gene expression analysis identified 10 clusters with distinct phenotypes and clearly distinguished between the two tissues (Figure 4C-E). The Ccr7.2 and Plasma clusters were observed solely in tumors, the Ccr7.1 cluster observed in both tumor and spleen, while the other cell phenotypes were primarily seen in spleens.

The Ccr7 clusters were characterized by expression of *Ccr7*, *Nr4a1*, *Junb*, *Cd83* and *Cd69* indicating an activated B cell state^37, 38^ (Figure 4C). Interestingly, the tumor-associated Ccr7.2 cluster, unique to the envRV vaccine, exhibited higher expression of these genes as well as clonal expansion, suggesting an increased activation level and migratory abilities compared to the Ccr7.1 cluster (Figure 4D-H). Importantly, the Plasma cluster, expressing the characteristic immunoglobulin components *Jchain*, *Iglv*, *Igkv*, and *Igkc,* was only found in tumors of envRV vaccinated animals, indicating a more differentiated cell state compared to the control groups (Figure 4F, H). In addition, the most expanded clones were found in plasma cell clusters as well as in the envRV group (Figure 4H, right panel and Figure 4I). In comparison to the tumor, the clonal expansion in the spleen was more limited and no statistical differences in cluster composition were observed between treatment groups (Figure S4B-C).

Given the evidence that the envRV RNA-LPX vaccine promotes the differentiation of B cells into plasma cells, we aimed to determine whether mutation-specific antibodies were generated upon vaccination. We collected serum from vaccinated tumor-bearing mice and tested it in an ELISA assay against the mutated full-length protein (474 AA) expressed by E0771 cells. In contrast to controls, mostly IgG antibodies were detected in the serum from envRV-treated animals (Figure 5A and S5A). To narrow down the recognition of serum-derived IgG antibodies to the mutated site of the full protein, we tested the envRV serum against the 29 AA-long mutated sequence encoded in the vaccine and found a similar titer curve (Figure 5A). These antibodies were predominantly of the IgG1 and IgG2c subclasses (Figure S5B).

**Figure 5:**
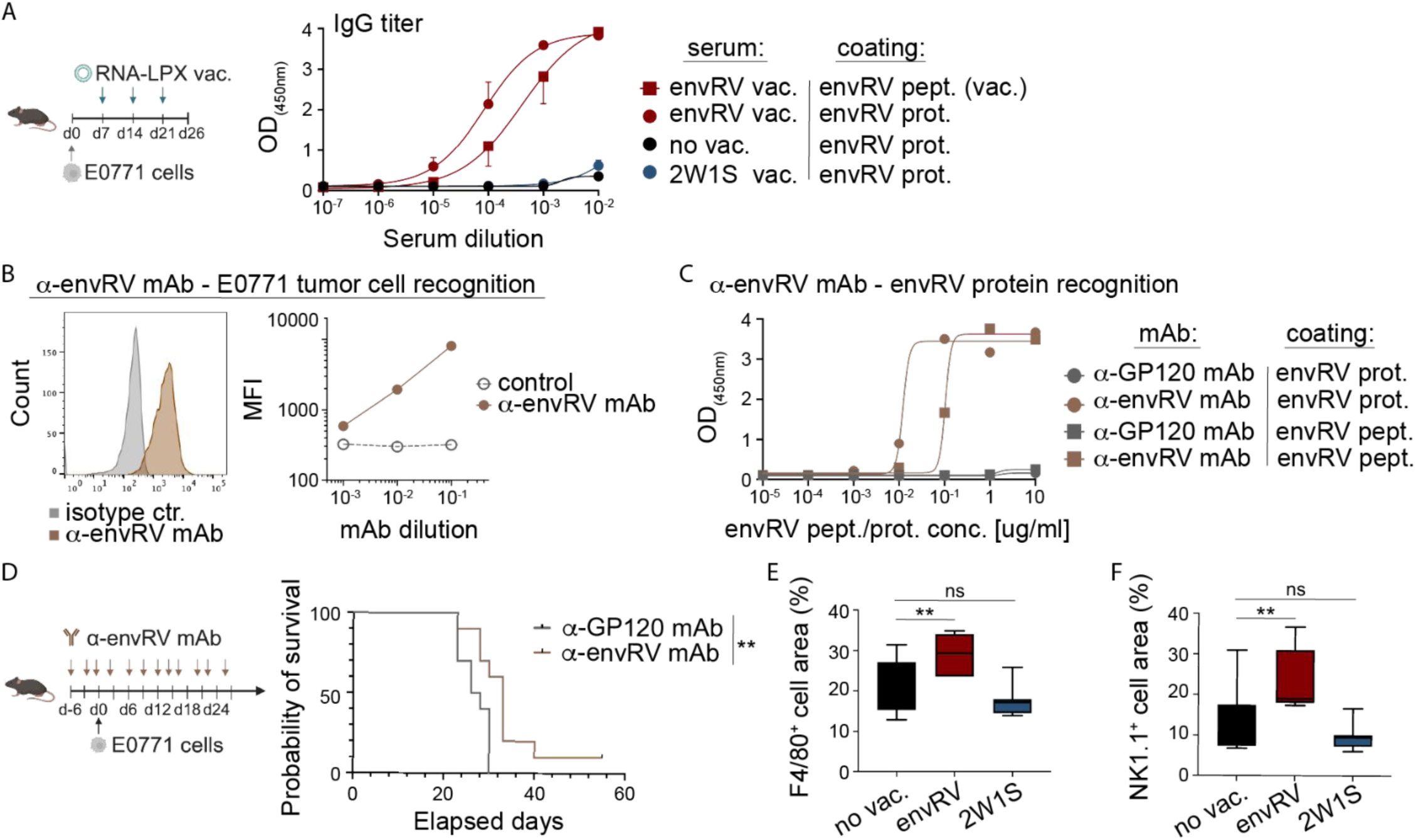
EnvRV RNA-LPX vaccine induces mutation-specific antibodies that prolong survival of mice bearing E0771 tumors. A: IgG titer determined by ELISA in serum from envRV-vaccinated mice recognizing the mutated full envRV protein (envRV prot.; red circle curve) or the mutated envRV peptide encoded in RNA-LPX vaccine (envRV pept. (vac.); red square curve). Serum from non-vaccinated (black curve) or 2W1S RNA-LPX vaccinated animals (blue curve) were used as control. Data are shown as (OD_450nm_) values for dilution of serum from 10^-2^ to 10^-7^. B: Flow cytometry plot and quantification graph showing the staining of the envRV-specific mouse monoclonal antibody (ɑ-envRV mAb) against E0771 tumor cells. Data are represented as MFI for three mAb dilutions (10^-1^ to 10^-3^). C: ELISA assay measuring recognition of envRV-specific mouse monoclonal IgG2a antibody (ɑ-envRV mAb) against the full envRV protein (envRV prot.; brown circle curve) or the mutated envRV peptide encoded in the RNA-LPX vaccine (envRV pept.; brown square circle). Mouse IgG2a GP120 monoclonal Ab was used as a control for each condition (grey lines). Data are shown as (OD_450nm_) values for various concentrations of coated envRV peptide or protein (10^-5^ to 10 μg/ml). D: Survival curve of E0771 tumor-bearing mice receiving envRV-specific mouse IgG2a (ɑ-envRV mAb) monoclonal antibody (brown line) or isotype control mIgG2a GP120 mAb (grey line). Data are shown as probability of survival for each elapsed day. **p<0.01 (Log-rank test). E-F: IHC staining for F4/80 and NK1.1 positive cells from tumors of non-vaccinated and envRV or 2W1S RNA-LPX vaccinated animals. Data are shown as boxplots with median of percentage of positive cell areas. ns, non-significant; **p<0.01.

To determine if there was a functional role for the envRV vaccine-induced antibodies, we sorted envRV-specific B cells from tumor-bearing mice and generated an ɑ-envRV specific monoclonal antibody (mAb) (see Methods). After confirming the specificity of the ɑ-envRV mAb towards E0771 tumor cells (Figure 5B) and the envRV protein (Figure 5C), we evaluated its anti-tumor activity *in vivo*. The survival of mice receiving the ɑ-envRV mAb was partly but significantly improved compared to those in the control group, suggesting that antibodies recognizing the mutated envRV protein promoted anti-tumor activity (Figure 5D).

Tumor-specific antibodies can eliminate tumor cells through various mechanisms, including antibody-dependent cellular cytotoxicity (ADCC) via NK cells and antibody-dependent cellular phagocytosis (ADCP) via macrophages. Interestingly, we observed a significant increase in intratumoral macrophages and NK cells in tumor samples from mice vaccinated with envRV compared to controls (Figure 5E-F).

### EnvRV RNA-LPX vaccine enhances endogenous intratumoral CD8^+^ T cell responses that control tumor growth

The vaccine-elicited Th1 response could conceivably have promoted CD8^+^ T cell responses leading to anti-tumor activity. To assess this possibility, we depleted CD8^+^ T cells in envRV vaccinated mice bearing tumors and found nearly complete loss of tumor control as compared to control (Figure 6A). Thus, CD8^+^ T cells were required for vaccine-induced anti-tumor activity despite the fact that the RNA-LPX vaccine did not contain any MHCI-restricted CD8^+^ T cell epitopes.

**Figure 6:**
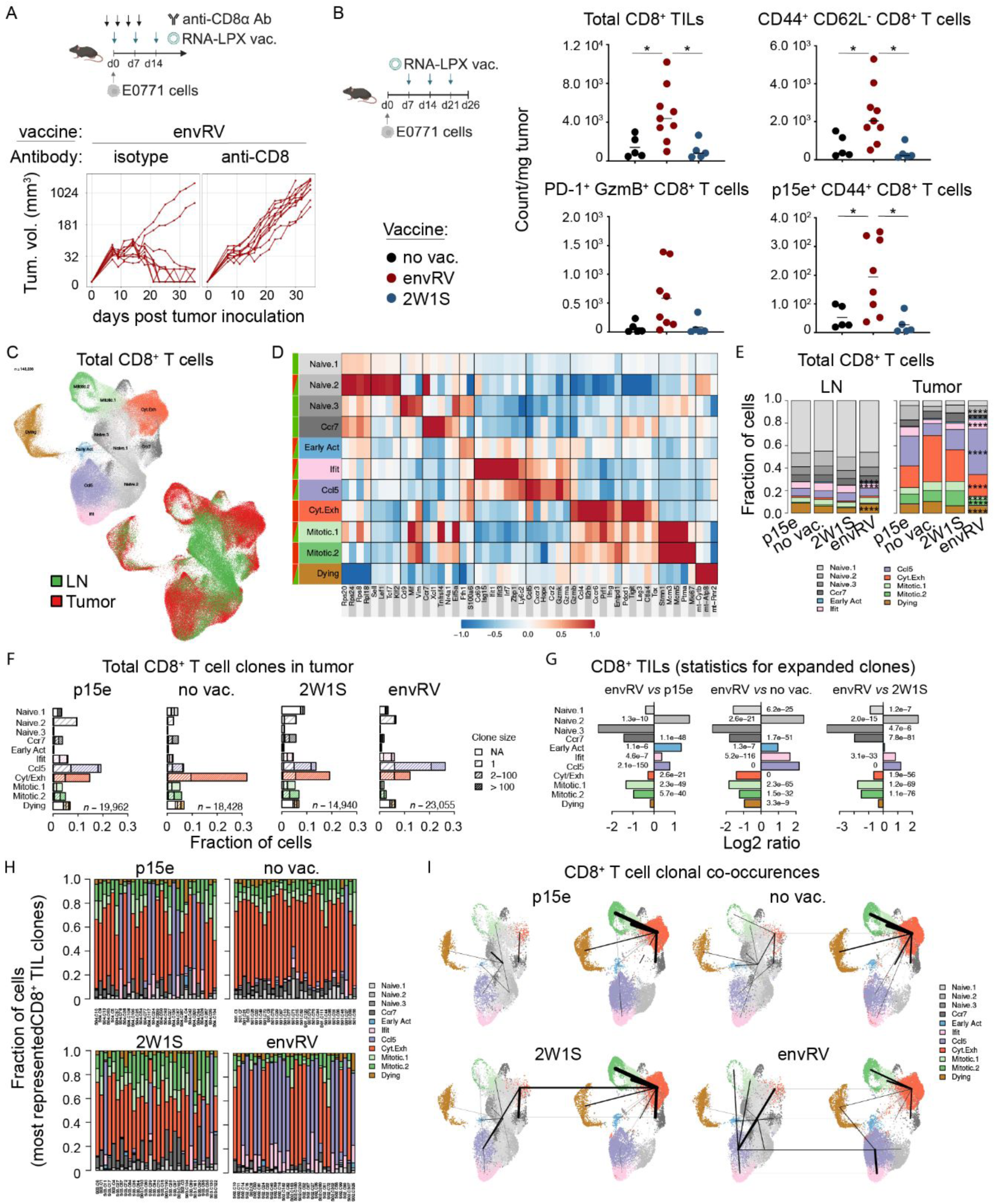
EnvRV RNA-LPX vaccine expands the endogenous progenitor-like CD8^+^ TILs responsible for tumor rejection. A: Study design and tumor growth curve of mice vaccinated with envRV RNA-LPX and treated with anti-CD8ɑ antibody or isotype control (20 mg/kg), received IV two days before tumor implantation then IP on days 1, 4, and 8. Tumor volumes were measured for 35 days post tumor challenge. B: Flow cytometry data summary of the mean count of total, effector memory (CD44^+^CD62L^-^), PD-1^+^Granzyme B (GzmB)^+^ and p15e^+^CD44^+^ CD8^+^ TILs per mg of tumor from mice vaccinated with envRV or 2W1S RNA-LPX vaccine. No vaccine was used as a control. Representative experiment out of three. *p < 0.05. C: UMAP of CD8^+^ T cells sorted from tumors and tdLNs clustered from scRNAseq data for all CD8^+^ T cells (left) and for each tissue (right). All groups were combined, n=143236 cells. See Figure S6D. D: Heatmap of relative average expression of selected marker genes associated with CD8^+^ T cell phenotype, function or differentiation states in each cluster identified in the UMAP in C. Tissues are represented by the color bar on the left (LNs, green; tumors, red). mt-:mitochondrial. E: Stacked bar graphs of CD8^+^ T cell cluster composition of all cells in LN and tumor from non-vaccinated, 2W1S (control MHCII), envRV (MHCII) or p15e (MHCI) RNA-LPX vaccinated mice. Statistical significance between envRV and all other groups are shown. *\*\*\*\**p<0.0001. F: Barplots showing the fraction of CD8^+^ T cells for each cluster identified from the UMAP in C, for the tumor tissue. In each stacked bar, open bar denotes cells for which TCR were unidentified, light stripes for singletons, medium stripes for clones between 2-100 cells, and high-density stripes for clones greater than 100 cells. Stripe color represents the primary cell phenotype of each clone as shown in the UMAP in C. G: Barplots showing the log2 ratio of the clonal expanded fraction of CD8^+^ T cells for each phenotypic cluster in tumors. Comparisons between envRV and each of the 3 other treated groups with significant statistical values are shown. H: Proportion of CD8^+^ T cell clusters for each of the 30 most represented CD8^+^ T cell clones in the tumor and for each treatment group. I: Cluster co-occurrence analysis in LNs and tumors from the no vaccine or the p15e, 2W1S, and envRV RNA-LPX vaccinated groups. Lines within UMAPs denote co-occurrence between different clusters. Thickness of line denotes relative strength of co-occurrence, with thickest lines indicating strongest co-occurrence.

We next assessed CD8^+^ T cell infiltration and expansion in tumors and tumor-draining lymph nodes (tdLNs) as tumor-specific CD8^+^ T cells are known to be mainly primed in tdLNs. We observed an accumulation of CD8^+^ T cells in both tissues, and an increased number of effector memory (CD44^+^CD62L^-^) and cytotoxic (PD-1^+^GzmB^+^) CD8^+^ TILs following the envRV vaccine (Figures 6B and S6B). In line with these results, serum from envRV vaccinated animals exhibited higher levels of CCL5, CXCL9 and CXCL10 chemokines compared to controls, which have been associated with increased T cell infiltration in tumor^39, 40^. In addition, envRV induced higher serum levels of pro-inflammatory cytokines (IL-12p40 and TNFɑ), with IL-12 having been associated with increased Th1 differentiation and CD8^+^ T cell clonal expansion^41, 42^, supporting a role for the vaccine-induced Th1 cells in promoting CD8^+^ T cell responses. Since enhanced cytokine/chemokine production was not observed in naive mice (not shown), the tumor is likely the tissue source. Importantly, the envRV vaccine increased tumor-specific CD8^+^ T cells in tumors, demonstrated by p15e-specificity; p15e is an MHCI-restricted epitope contained within the MuLV envelope protein and distinct from the envRV MHCII-restricted epitope^43^ (Figure 6B). These p15e-specific CD8^+^ T cells can recognize and eliminate E0771 cancer cells following p15e RNA-LPX vaccination in a prophylactic setting (Figure S6C).

We next characterized the nature of the envRV-expanded CD8^+^ T cells from tdLNs and tumors using scRNAseq and scTCRseq (Figure S6D). Mice vaccinated with the p15e RNA-LPX were also included in the analysis.

Unsupervised clustering analysis revealed that at steady state there were marked differences between CD8^+^ T cells in tumors versus tdLNs, with some overlapping clusters (Figure 6C-D). We investigated how these CD8^+^ T cell subtypes were affected by the envRV vaccine in both tissues. The most prevalent clusters in the tdLNs did not exhibit differences in their proportions across the treatment groups (Figure 6E). However, the envRV vaccine decreased the proportion of the Ifit cluster showing high expression of the early activation molecule *CD69*, the IFN-associated genes *Irf7, Isg15, Ifit1* and *Ifit3*, and the cytotoxic molecules *Gzmk*, *Gzma* and *Gzmb* compared to other treatment groups (Figures 6D-E).

Within the tumor, the envRV group showed a decreased fraction of the Ccr7 cluster that upregulated the transcripts associated with T cell exhaustion (*Tox*, *Pdcd1*, *Lag3)* as well as the two mitotic clusters expressing primarily markers of mitosis and inhibitory molecules (Figures 6D-F). These results indicated that there was a lower level of exhaustion in envRV-expanded CD8^+^ TILs. In parallel, the envRV vaccine increased the proportion of the Ccl5 cluster in the tumor compared to all other groups (Figure 6E-F). This cluster highly expresses the chemokine *Ccl5,* the chemokine receptors *Cxcr3 and Ccr2*, and the activation/cytotoxic molecules *Cd69, Gzmk and Gzma*.

In contrast to its LN counterpart, the Ifit cluster increased in tumor of envRV-treated mice compared to non-treated and 2W1S groups (Figure S6E). Interestingly, both Ifit and Ccl5 clusters expressed higher levels of molecules associated with stem-like/progenitor T cell states (*Tcf7*, *Lef1*) than other non-naive CD8^+^ TIL clusters (Figure S6G).

In addition, the envRV vaccine group contained fewer proportions of total CD8^+^ Cytotoxic/Exhausted (Cyt/Exh) TILs that upregulated genes encoding cytotoxic molecules (*Gzmb, Prf1, Ifng*) as well as inhibitory molecules (*Ctla4, Tox, Pdcd1, Tigit, Lag3*) compared to untreated and 2W1S groups (Figure S6E). These results suggest that envRV-mediated CD8^+^ T cell accumulation may have been more recently primed, with fewer exhausted and more activated CD8^+^ TILs.

We then analyzed the degree of clonal expansion of CD8^+^ T cells within each cluster using scTCRseq. While no T cell clonal expansion was detected in dLNs (Figure S6F), the Ccl5 and Cyt/Exh clusters in tumors contained the highest proportions of expanded clones across samples (Figure 6F-G). The Cyt/Exh cluster derived primarily from hyper-expanded clones (clone size >100) across all groups. Interestingly, the envRV group showed a significantly higher clonal expansion of Ccl5 CD8^+^ TILs, including increased number of hyper-expanded clones (>100), as well as a reduced proportion of Cyt/Exh hyper-expanded clones in comparison to all control groups (Figure 6F-G). This result aligns with our recent findings^44^, which demonstrated that efficacious immunotherapy combining PD-L1 and TIGIT blockade specifically expands Ccl5 CD8^+^ TILs in CT26 murine tumors. In addition, the 30 largest CD8^+^ T cell clonotypes in tumors from untreated and 2W1S vaccinated mice were composed primarily of cells with a Cyt/Exh transcriptional profile in contrast to the major Ccl5 phenotype found in the envRV group (Figure 6H). Importantly, while the expanded Cyt/Exh clones were significantly reduced in the envRV group compared to p15e (Figure 6G), the Ccl5 phenotype was increased in the most represented clones following envRV vaccine compared to p15e RNA-LPX (Figure 6H). Together, these results indicate that the envRV vaccine shifted the intratumoral CD8^+^ T cell population from a more terminally differentiated or exhausted Cyt/Exh phenotype to a “younger” Ccl5 phenotype. Thus, the MHCII vaccine appeared to expand cells with progenitor/stem-like and effector characteristics, even more effectively than the MHCI p15e vaccine.

To investigate in greater detail the origin and trajectories of individual CD8^+^ TCR clonotypes, we analyzed clonal co-occurrences among CD8^+^ T cell clusters and tissues using TCRseq (see Methods). Within the untreated and 2W1S groups, expanded CD8^+^ T cell clones largely had co-occurrences involving the Cyt/Exh cluster (Figure 6I). In contrast, expanded CD8^+^ T cell clonotypes from envRV vaccinated mice tended to have additional co-occurrences involving the Ccl5 cluster. Interestingly, even though the p15e vaccine also showed Ccl5 CD8^+^ TIL expansion, the main clonal co-occurences still involved the Cyt/Exh cluster (Figure 6E-G, I). These results suggest that, unlike the p15e vaccine, the envRV vaccine not only boosted the number of Ccl5 CD8^+^ T cell clones but also enhanced the quality of CD8^+^ TIL responses. These data are consistent with the shift in clonal expansion and phenotypes toward the Ccl5 cluster by the envRV vaccine.

Considering the differences in phenotypic transcripts and clonal expansion induced by envRV and the MHCI p15e vaccines, we explored whether these variations would lead to different anti-tumor responses in the context of established tumors. Indeed, tumor-bearing mice vaccinated with p15e RNA-LPX did not reject established tumors, unlike those treated with the envRV vaccine (Figure S6H). This suggests that the ability of the MHCII envRV vaccine to produce a less terminally differentiated CD8^+^ T cell phenotype may partially explain the greater efficacy seen with this approach.

### EnvRV-enhanced CD8^+^ TIL activity is dependent on cDC1s and tumor-specific CD4^+^ T cells at the tumor site

cDC1s and CD4^+^ T cells are critical components for eliciting and maintaining cytotoxic CD8^+^ T cell responses in a context of tumor^45^.

Considering the reduction in vaccine-induced CD4^+^ T cells in cDC1-depleted mice, we next assessed the impact of cDC1 deletion on the envRV-enhanced CD8^+^ response. As expected given the critical role of cDC1 in priming CD8^+^ T cells^46^, there was a net reduction in the number of total CD8^+^ T cells in the tumors from *Batf3^-/-^* mice, which mostly contain antigen experienced T cells (Figure S7A). In contrast, the proportion of CD8^+^ T cells in spleen and tdLN, which contained a large fraction of naive T cells, was unaffected by the absence of cDC1 (Figure S7A).

Indeed, the frequency of PD-1^+^Gzmb^+^ cytotoxic CD8^+^ TILs was lower in *Batf3^-/-^* compared to WT mice, consistent with reduced effector differentiation of the few primed TILs (Figure S7B).

The reduction of CD8^+^ TILs in envRV-vaccinated *Batf3^-/-^*mice likely reflected the combination of deficient priming or boosting by cDC1 combined with reduced envRV-specific CD4 help. To determine if the cDC requirement was restricted to the priming phase, we developed a conditional knockout model using a Diphtheria toxin (DT)-induced cDC conditional deletion. DT treatment in CD11c.cre^+/-^ *Zbtb46*.LSL.DTR^wt/loxp^ (CD11c-DTR) chimeras, efficiently depleted cDCs up to 48h post treatment initiation (Figure S7C). DT could thus be used to selectively deplete cDCs in tumor-bearing chimeric mice during vaccine priming and/or boosting. As shown in Figure 7A, the proportion of CD8^+^ TILs was significantly lower when CD11c^+^ DCs were depleted during envRV priming or boosting, indicating that cDCs are necessary for both the envRV-vaccine priming and boosting phases.

**Figure 7:**
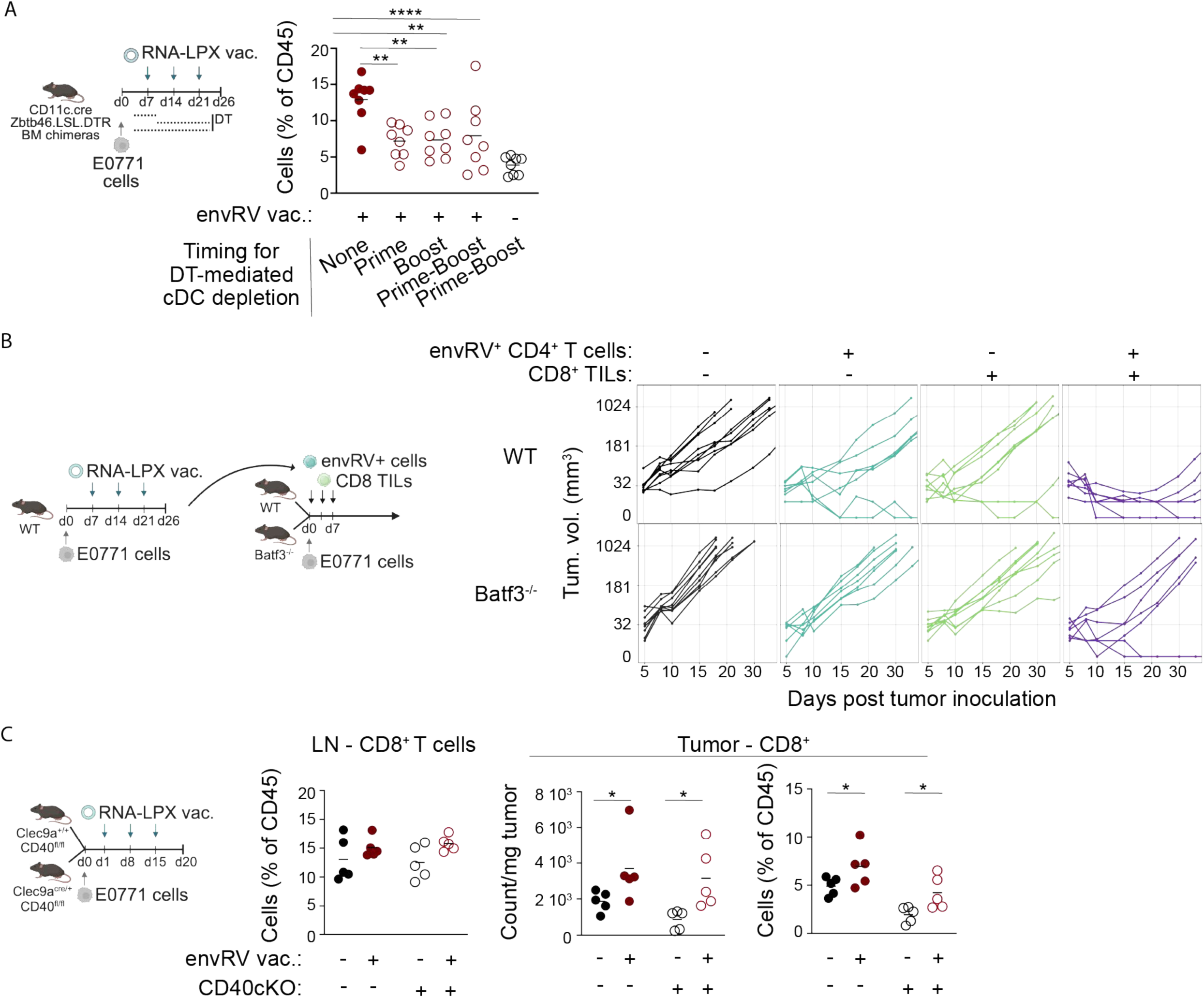
EnvRV RNA-LPX-mediated effector activity of CD8^+^ TILs depends on cDC1s and partially on CD40 expression in cDCs. A: Proportion of CD8^+^ TILs among CD45^+^ cells from tumor-bearing CD11c.cre ZBTB46.LSL.DTR bone marrow chimeras vaccinated with envRV RNA-LPX and treated or not with diphtheria toxin (DT) at the time of priming and/or boosting. Data are shown as mean. **p<0.01, ****p<0.0001. See also Figure S7B. B: Adoptive cell transfer study. Tumor-bearing donor WT mice were vaccinated with envRV RNA-LPX vaccine. Tumors and spleens were harvested 5 days after the third vaccination. EnvRV^+^ CD4^+^ T cells sorted from spleens and tumors were pooled. CD8^+^ T cells sorted from tumors were pooled. Cells were intratumorally transferred into WT or Batf3^-/-^ tumor-bearing recipient animals either separately or combined on days 1, 4 and 7 post tumor inoculation. PBS injection was used as control. Tumor growth curves of recipient mice are shown for each treatment group. C: Clec9a^cre/+^CD40^fl/fl^ (CD40 cKO) and Clec9a^+/+^CD40^fl/fl^ littermate (control) mice were vaccinated or not with envRV RNA-LPX, 7 days apart starting one day post E0771 tumor inoculation. Tumors and tdLNs were harvested 5 days post last vaccine for flow cytometry analysis. For LNs, frequency of CD8^+^ T cells among CD45^+^ cells is shown. For tumors, count of CD8^+^ TILs and proportions of CD8^+^ TILs among CD45^+^ cells are shown. Data are shown as mean.*p<0.05. n=1 experiment.

To determine whether envRV-specific CD4^+^ TILs and cDC1s were required to maintain the CD8^+^ TIL effector activity, we took an approach recently published by Sultan et al.^24^ using CD4^+^ and CD8^+^ T cell intratumor adoptive transfer. We adoptively transferred envRV-specific CD4^+^ T cells from spleen and tumor or CD8^+^ TILs from previously envRV vaccinated tumor-bearing WT donor mice into tumors of untreated WT and *Batf3^-/-^* recipient mice (Figure 7B). While intra-tumoral transfer of envRV-specific CD4^+^ T cells alone slightly delayed tumor growth, consistent with improved endogenous CD8^+^ T cell responses in WT animals, the transfer of CD8^+^ TILs alone had no effect on tumor growth. In contrast, only the co-transfer of both envRV-specific CD4^+^ T cells and CD8^+^ TILs efficiently controlled the tumor in WT animals. Importantly, in cDC1-deficient mice, none of the T cell transfers controlled tumor growth (Figure 7B). These results demonstrate that there is also a requirement of cDC1s at the tumor site for promoting effective anti-tumor activity of CD4^+^ and CD8^+^ T cells.

Finally, we assessed whether CD40/CD40L signaling was required for envRV-specific CD4^+^ T cells to provide help to cDC1s for clonal expansion of CD8^+^ T cells, using CD40 cKO mice. CD40 deficiency in cDCs did not impact the frequencies of CD8^+^ T cells either in tdLNs or in tumors, even following therapy (Figure 7C), suggesting that CD40 signaling in cDCs is not required for envRV-mediated CD8^+^ TIL expansion.

## Discussion

It has long been known that CD4^+^ T cells are critical for the generation of CD8^+^ T cell responses by providing “T cell help”, which is usually understood to reflect the ability of CD4^+^ T cells to provide cytokine or other signals directly to CD8^+^ T cells or indirectly by licensing DCs to present MHCI-bound antigens. In the context of therapeutic cancer vaccines, several studies have suggested that eliciting CD4^+^ T cells alone may be sufficient to generate anti-tumor responses^6, 47, 48^. Conceivably, cytotoxic cytokines or direct killing of MHCII-positive tumor cells might explain an inherent anti-tumor activity. Our work was intended to take a detailed look into the nature of the vaccine-induced CD4^+^ T cell response, by devising an approach that allowed us to investigate the effects of a well-defined and selective MHCII-dependent CD4^+^ T cell response. We have shown that vaccinating against the envRV antigen expressed by E0771 tumors orchestrated a multifaceted CD4^+^ T cell-dependent response involving the induction of not only Th1 cells but also Tfh cells that, in turn, elicited anti-tumor antibodies and cytokines that contributed to tumor elimination. More surprisingly, however, was the ability of this response to generate potent CD8^+^ T cells against at least one endogenous tumor antigen in a fashion dependent not only on CD4^+^ T cells but also cDC1s. This complexity may explain the ability of the RNA-LPX vaccine to generate (via Tfh cells) the IL-21-dependent production of anti-tumor antibodies as well as an endogenous CD8^+^ T cell response, which may be supported at least in part by Th1-derived IFN𝛾-mediated activation of DCs and macrophages.

We also observed the virtual elimination of neoantigen-specific Tregs at the tumor site following envRV RNA-LPX vaccine. Although decreases in total Tregs in tumors after mRNA vaccination have been observed previously^6^, the use of antigen-specific MHCII tetramers allowed us to conclude that this effect was specific to the relevant neoantigen-specific Treg population. The question of whether the vaccine drives Treg conversion to an effector phenotype or if they are simply diluted out by effector cells remains to be investigated.

Our results indicate that vaccinating with a short single MHCII neoepitope can provide the CD4^+^ T cell co-stimulation needed for B cell differentiation. Beyond promoting Th1 cells and enhancing tumor-specific CD8^+^ T cell responses, the MHCII mRNA-LPX vaccine also significantly expanded Tfh cells. This induction of Tfh differentiation - alongside IL-21 signaling - both of which are indispensable to B-cell maturation^49–52^ - appeared to drive the heightened activation state of B cells, the increase in memory B cell frequency, and the clonal expansion of plasma B cells. Together, these events facilitated a role for B cells in MHCII RNA-LPX vaccine-induced efficacy, namely the production of neoantigen-specific antibodies and their role in anti-tumor immunity. As B cells may also directly kill tumor cells^53, 54^, further studies will be required to evaluate the contribution of each mechanism independently.

While spontaneous antibody responses against tumor-associated antigens are frequently detected in the serum of cancer patients, their contribution in anti-tumor immunity remains unclear, especially in terms of clinical response^55^. Here, we observed the production of neoantigen-specific antibodies that did appear to exhibit anti-tumor efficacy, presumably by enhancing antibody-dependent cellular cytotoxicity (ADCC). Interestingly, we observed an intratumoral increase of both NK cells and macrophages following the neoantigen-specific vaccine, suggesting their involvement in vaccine-induced efficacy. In addition, through antigen-antibody complexes that can activate Fc𝛾R-mediated endocytosis, these vaccine-induced antibodies may promote DC maturation, thereby enhancing antigen presentation to T cells^56, 57^.

Perhaps the most striking result was the finding that an mRNA vaccine expressing a single MHCII-restricted neoantigen can provide sufficient help to endogenous CD8^+^ T cells to generate a protective response without the need to concurrently include an MHCI neoantigen. Importantly, the CD4^+^ T cells must recognize a tumor-associated neoantigen as generating CD4^+^ T cell responses to a control antigen did not produce an anti-tumor CD8^+^ T cell response. Our results clearly demonstrate that vaccine-elicited CD4^+^ T cell responses can achieve a form of epitope spreading that yields endogenously elicited CD8^+^ T cell responses. This is not always the case, however, as prior work has suggested that CD4^+^ T cell responses alone can facilitate effective tumor immunity following immunization with MHCII-restricted epitopes using RNA-LPX^6–8^. The mechanism of tumor killing has not been established but may reflect the B cell-dependent responses characterized here, or direct killing of MHCII^+^ tumor cells by cytotoxic CD4^+^ T cells. Direct killing by CD4^+^ T cells was not relevant in our studies since MHCII-KO E0771 tumors were eradicated as well as controls. As a result, when CD8^+^ T cells are involved, we were able to confirm a dependency on cDC1s in both dLN and tumor. Deletion of cDC1s demonstrated their role not only to support the vaccine response, but also following transfer of both envRV^+^ CD4^+^ T cells and CD8^+^ TILs from vaccinated WT tumor-bearing mice into Batf3^-/-^ animals. cDC1s have been shown to prime cognate CD4^+^ T cells and CD4^+^-helped cDC1s provide necessary support for CD8^+^ T cell anti-tumor responses^17^. Similarly to Ferris et al^17^, we found that cDC1s primarily prime CD4^+^ T cells, even in the context of RNA-LPX vaccination. Surprisingly, while CD40 signaling was required in the spleen to generate vaccine-specific CD4^+^ T cells, lack of CD40 in cDCs did not impair their accumulation in the tumor. Intratumoral cDC1s are also known to be essential to sustain anti-tumor CD8^+^ T cell responses^40, 58–60^. In line with recent reports showing the existence of intratumoral “triads” comprising DCs, CD4^+^ T cells and progenitors-like CD8^+^ T cells^20, 21^, we found higher expression of *Tcf7*, *Lef1*, and *GzmK* genes in envRV-expanded CD8^+^ TILs. However, accumulation of the TILs following MHCII vaccination was independent of CD40 signaling in cDCs. Thus, our results not only highlight the importance of cDC1s in priming vaccine-induced CD4^+^ T cells, but also support the crucial role of CD4^+^ T cell licensing of cDC1s for CD8^+^ T cell-driven tumor rejection in the context of cancer vaccination. Additional studies will define the precise mechanisms for cDC and CD4^+^ T cell help of CD8^+^ TILs for generating efficient tumor elimination.

MHCII-restricted neoantigen RNA-LPX redirects the clonal expansion toward progenitor-like CD8^+^ T cells in tumors. Strikingly, envRV-expanded CD8^+^ TILs resemble the multipotent CD8^+^ T cell clones we recently identified^44^, which significantly expanded within tumors following anti-PD-L1 and anti-TIGIT combination therapy. Interestingly, the MHCII-restricted mRNA vaccine was superior to the RNA-LPX vaccine encoding the p15e MHCI-restricted with respect to generating “young” or progenitor-like CD8^+^ T cells.

In conclusion, our study provides compelling evidence for the critical roles of CD4^+^ T cells, cDC1s and CD8^+^ T cells in orchestrating a comprehensive and effective anti-tumor immune response following MHCII neoantigen-restricted RNA-LPX vaccination. These findings have obvious implications for understanding the role and significance of CD4^+^ T cell responses when designing cancer vaccines, especially since eliciting this class of response is typically associated with the generation of antibody-based humoral immunity for prophylaxis. It remains to be seen whether it will always be the case that CD8^+^ T cells expanded following an RNA-LPX encoded MHCII vaccine are always superior to CD8^+^ T cells expanded by MHCI-restricted vaccination. One possible explanation may lie in the relative routes of antigen delivery. In our experiments, the RNA-LPX was administered intravenously, meaning that the initial priming of T cell immunity likely occurred in the spleen. From there, the neoantigen-specific CD4^+^ T cells would likely traffic to dLN and the tumor, where they would provide help to local cDC1s displaying MHCI-restricted neoepitopes following the cross-presentation of tumor neoantigens. Conceivably, the spatial separation of these two events may contribute to a more robust and effective CD8^+^ T cell response.

## Methods

### Mice

Animals were maintained in accordance with the Guide for the Care and Use of Laboratory Animals (National Research Council 2011). Genentech is an AAALAC-accredited facility and all animal activities in this research study were conducted under protocols approved by the Genentech Institutional Animal Care and Use Committee (IACUC).

C57BL/6N mice were purchased from Charles River Labs (Hollister, CA) or SLAC Laboratory Animal Co., LTD. C57BL/6J mice were purchased from the Jackson Laboratory (JAX stock #000664). IL21R KO mice were purchased from the Jackson Laboratory (B6.129-*Il21rtm1Kopf*/J, JAX stock #019115). Batf3 KO mice were purchased from the Jackson Laboratory (B6.129S(C)-*Batf3tm1Kmm*/J, JAX stock #013755) or bred in house (BATF3.exon1exon2.noneo.ko.B6J) on a BL/6J background. CD40 conditional KO mice (CD40.CRISPR.flox.cko_CLEC9A.iCre.noneo.ki - MNV.Helico Free) were obtained and bred in house on a BL/6 background by crossing CLEC9A.iCre.noneo.ki - MNV.Helico Free mice bred in house and C57BL/6JGpt-Cd40em1Cflox/Gpt mice purchased from GemPharmatech (Strain NO.T052302). For bone marrow transfer experiments, CD45.1 C57BL/6J recipient mice (B6.SJL-Ptprca Pepcb/BoyJ, JAX stock #002014) were lethally irradiated twice with 500 RAD for 4 minutes with a 3-hour interval in between irradiation before being reconstituted the day after with 4.5-7.2 million bone marrow cells isolated from ZBTB46.LSL.DTR.mCherry.IRES.cki_CD11c.Cre.tg CD45.2 donor mice (crossed and bred in-house). Diphtheria toxin (DT) was diluted in PBS and injected intraperitoneally during RNA-LPX vaccine prime; on day 6 post tumor implantation (500 ng/mouse) and days 7, 8, 9 and 10 (100 ng/mouse) or during vaccine boosts; on day 13 (500 ng/mouse) and days 14, 15, 16 and 17 (100 ng/mouse) and on day 20 (500 ng/mouse) and days 21, 22, 23 and 24 (100 ng/mouse). Females 6-16 weeks of age were used for experiments. All mice that were housed at Genentech were in individually ventilated cages within animal rooms maintained on a 14:10h, light:dark cycle. Animal rooms were temperature and humidity-controlled, between 20 and 26°C and 30 and 70% humidity, respectively, with 10 to 15 room air exchanges per hour. Mice were acclimated to study conditions for at least 3 days before tumor cell implantation. All animal studies were approved by Genentech’s Institutional Animal Care and Use Committee and adhere to the NRC Guidelines for the Care and Use of Laboratory Animals.

### Cell Lines

The E0771 cell line was purchased from CH3 Biosystems. E0771 MHCII KO were generated by subsequently knocking out H-2Aa (sgRNA H-2Aa 42F + H-2Aa 46F) and H-2Ab (sgRNA H-2Ab 06R + H2-Ab 59F) genes using the CRISPR KO methodology using the program CA137 and buffer SE on the Amaxa transfection system (Lonza). Transfected cells were maintained in culture and stimulated with IFN-γ at 20 ng/ml for 72 hours to confirm the absence of MHCII expression. E0771 MHCII KO cells were sorted by FACS to obtain a pure E0771 MHCII KO population. All cell lines were stored by a common cell repository at Genentech. Cell lines are routinely screened, and cells used in this study were negative for mycoplasma and authenticated by RNAseq analysis. Two methods of mycoplasma detection were used to avoid false positive/negative results: Lonza Mycoalert and Stratagene Mycosensor. Cells were cultured in RPMI 1640 medium plus 2 mM L-glutamine with 10% fetal bovine serum (FBS; Hyclone, Waltham, MA). Cells in log-phase growth were centrifuged, washed with Hank’s balanced salt solution (HBSS), counted, and resuspended in 50% HBSS and 50% Matrigel (BD Biosciences; San Jose, CA).

### RNA-LPX vaccines

mRNAs encoding for the envRV neoantigen (FSPPPGPPCCSGSRVSTPGCSRDCEEPLT) or 2W1S MHCII-restricted antigen (EAWGALANWAVDSA) or for the p15e tumor-associated MHCI restricted antigen (KSPWFTTL) were synthesized by Genentech. RNA was formulated with liposomes consisting of DOTMA and DOPE at a charge ratio (+):(-) of 1.3:2, yielding negatively charged RNA-LPX.

### *In vivo* tumor studies

Female mice were housed at Genentech in standard rodent microisolator cages, and acclimated for at least 3 days before cell injection. One efficacy study was outsourced to WuXi (China). E0771 cells were inoculated in the left #4 or #5 mammary fat pad (1x10^5^ cells in 100 μl of HBSS/Matrigel 1:1 mixture). When indicated mice were treated with neutralizing antibodies for mouse IFN-γ, mouse CD8, mouse CD20, or corresponding isotype control antibodies. Animals were treated i.v. with envRV RNA-LPX vaccine or 2W1S RNA-LPX vaccine on days 7, 14 and 21 or otherwise indicated. For pharmacodynamic/phenotyping studies, mice were euthanized 5 days after the last vaccination and spleens, LN and tumors were collected for flow cytometry analysis. For therapeutic efficacy studies, tumors were measured 2 times per week using digital calipers, and tumor volumes calculated using the modified ellipsoid formula, 1/2 x (length x width^2^). Mice were euthanized if tumors ulcerated or volumes exceeded 1500 mm^3^. Early euthanasia of some mice due to tumor ulceration contributed to some variability in final sample sizes of immune pharmacodynamic studies. No mice met criteria for euthanasia due to body weight loss or adverse clinical signs. Sample sizes in the mouse studies were based on the number of mice routinely needed to establish statistical significance based on variability within study arms. Investigators were not blinded to treatment.

### Antibodies for *in vivo* studies

*In vivo* IFN-γ depletion was performed by treating mice i.p. with 1 mg of anti-mouse IFN-γ (Bioxcell, BE0054, clone R4-6A2, Rat IgG1, κ), on day 1 after E0771 tumor implantation followed by 500 μg on day 2, 3, 4, 5 and 11. *In vivo* B cell depletion was performed by treating mice with anti-mouse CD20 (Genentech, 8596, Mouse IgG2a) 10 mg/kg i.v. the day before E0771 tumor implantation and 5 mg/kg i.p. on day 3, 6, 10, 13 and 17. *In vivo* CD8^+^ T cell depletion was performed by treating mice with anti-mouse CD8ɑ (Genentech, anti-CD8:9805, rat IgG2b) 20 mg/kg i.v two days before tumor implantation followed by 20mg/kg i.p. on day 1, 4, and 8. The rat IgG2a (Biocell, BE0090, clone LTF-2) was used as an isotype control.

### Tissue preparation and flow cytometry

E0771 tumors were collected, weighed, and digested using tumor dissociation kit solution (Miltenyi, 130-096-730) in GentleMACS C tubes (Miltenyi, 130-093-237) using the program mouse 37C_m_TDK_1. Spleens were collected in cold PBS and single-cell suspensions were generated by mashing the spleen tissue through a 70-μm cell strainer (BD Falcon, 08-771-19) in Hank’s-based Cell Dissociation Buffer (Gibco, 13-150-016) supplemented with Liberase TL (Roche, 05401020001) and DNase I (ThermoFisher Scientific, 9008). Red blood cells were lysed with ACK lysis buffer (in-house). Samples were then passed through a 70 µm filter and prepared for flow cytometry. Cells were transferred into a V bottom 96-well plate, washed twice with HBSS (Gibco, 14025-076), before being stained at room temperature with HBSS solution containing 5 μg/ml of Fc receptor blocking antibody (BD biosciences, 553142) and either with 20 μg/ml of pMHCI tetramer for 30 minutes or with 12.5 μg/ml of pMHCII tetramer for 1 hour. Tetramer reagents were generated in-house. Cells were then labeled for surface antigens with monoclonal antibodies together with LIVE/DEAD Fixable (ThermoFisher) at 4°C for 20 minutes. For intracellular staining of GzmB, cells were fixed with Cytofix/Cytoperm buffer (BD Biosciences, 554722) for 20 minutes at 4°C, washed twice with Perm/Wash buffer (BD Biosciences, 554723) and stained with intracellular antibodies diluted in Perm/Wash buffer for 1 hour at 4°C. For intranuclear staining of transcription factors, cells were fixed with Foxp3 / Transcription factor fixation buffer (eBioscience, 00-5523-00) for 1 hour in the dark at 4°C and then washed twice with perm/wash buffer (eBioscience, 00-8333-56) and stained with a mix of nuclear antibodies for 1 hour at 4°C in the dark. Appropriate isotype controls were included for staining of transcription factors. Cells were collected on a Symphony flow cytometer (BD Biosciences) and analyzed with FlowJo software (Version 10.8.1, Treestar, Ashland, OR, USA).

### Single-cell RNAseq and TCR V(D)J clonotype profiling

#### CD4^+^ T cells scRNA/TCR library preparation

E0771 tumor-bearing mice were vaccinated with envRV or 2W1S RNA-LPX on day 7, 14 and 21 and tissues were harvested on day 26. Cells from tumors and spleens were stained with PE- and APC-conjugated MHCII tetramers and enriched for CD4^+^ tetramer^+^ cells by positive selection using anti-PE (Miltenyi, 130-048-801) and anti-APC MicroBeads (Miltenyi, 130-090-855) and LS columns (Miltenyi, 130-042-401) following the manufacturer’s instructions. Enriched cells were subsequently stained for surface antibodies (Live/Dead, FcBlock, CD45, CD90.2, CD4, CD8, CD44, Dump (MHCII, CD11b, gdTCR, F4/80, CD19, NK1.1, CD11c)) and viable CD4^+^ T cells were sorted using a BD FACSAria™ Fusion flow cytometer (BD Biosciences, San Jose, CA). Separately sorted tetramer^pos^ CD44^+^ and tetramer^neg^ CD44^+/-^ CD4^+^ T cells were then mixed in equal proportion for further NGS analysis. For scRNA/TCRseq, 2,000–50,000 cells were sorted in 300 μL of PBS + 2% FCS and kept at 4 °C. Cells were then counted and resuspended at an adequate concentration for loading into the 10X chip. Gene expression and TCR libraries were generated using the Chromium Single Cell 5′ Library and V(D)J Reagent Kit (10X Genomics) according to the manufacturer’s recommendations. 20,000 cells per sample were loaded into each channel of the Chromium Chip, and recommendations were followed assuming targeted cell recovery of 2,000–10,000 cells. 10X Geomics single-cell gene expression libraries were quantified with the Qubit dsDNA HS Assay Kit (Thermo Fisher Scientific) and profiled using the Bioanalyzer High Sensitivity DNA Kit (Agilent Technologies). Gene expression libraries were sequenced on the NovaSeq 6000 (Illumina) to generate ∼200 million paired-end reads per library in a 28x10x10x90 bp configuration. TCR libraries were sequenced on NovaSeq 6000 (Illumina) to generate 40-50 million paired-end reads per library in a 28x10x10x90 bp configuration.

#### CD45^+^ and B220^+^ cells scRNA/ADT/TCR/BCR library preparation

E0771 tumor-bearing mice were left unvaccinated or vaccinated with envRV or 2W1S RNA-LPX on day 7, 14 and 21 and tissues were harvested on day 26. Cells from spleen were stained for surface antibodies (Live/Dead, CD45, CD90.2, CD4, CD8, B220, Dump (CD11b, F4/80, NK1.1, CD11c)), plus one of the following hashtag antibodies (TotalSeq™-C0301 (barcode sequence: ACCCACCAGTAAGAC), -C0302 (barcode sequence: GGTCGAGAGCATTCA), -C0303 (barcode sequence: CTTGCCGCATGTCAT), -C0305 (barcode sequence: CTTTGTCTTTGTGAG), -C0306 (barcode sequence: TATGCTGCCACGGTA), -C0307 (barcode sequence: GAGTCTGCCAGTATC), -C0308 (barcode sequence: TATAGAACGCCAGGC), -C0309 (barcode sequence: TGCCTATGAAACAAG), or -C0310 (barcode sequence: CCGATTGTAACAGAC). Following processing, tumor samples were enriched for live cells using the Dead Cell Removal Kit (Miltenyi, 130-090-101). Cells were then labelled with antibody-derived Tags (ADT) TotalSeq™-C0961 PE-conjugated (Biolegend, 405155) and TotalSeq™-C0957 APC-conjugated (Biolegend, 405285) MHCII tetramers and then stained for surface antigens (Live/Dead, CD45, CD90.2, CD4, CD8, B220, Dump (CD11b, F4/80, NK1.1, CD11c)) plus one of the following Hashtag antibodies for each mouse: TotalSeq™-C0301, -C0302, -C0303, -C0305, -C0306, -C0307, C0308, -C0309, or -C0310. Cells from spleen or tumor were pooled by treatment and viable CD45^+^ B220^+^ Dump^-^ CD4^-^ CD8^-^ CD90.2^-^ from spleen and CD45^+^ cells from tumor were sorted using a BD FACSAria™ Fusion flow cytometer (BD Biosciences, San Jose, CA). For scRNA/TCR/BCRseq, preparation of samples for loading to the 10X chip as well as reagents and instruments was performed as described in the CD4^+^ T cell library preparation section above.

#### CD8^+^ T cells scRNA/TCRseq library preparation

E0771 tumor-bearing mice were left unvaccinated or vaccinated with envRV, 2W1S or p15e RNA-LPX on day 7, 14, and 21 and tumors and dLNs were harvested on day 26. Following processing, tumor samples were enriched for live cells using the Dead Cell Removal Kit (Miltenyi, 130-090-101). Cells from tumor or dLN were then stained for surface antigens (Live/Dead, CD45, CD90.2, CD4, CD8, CD44, Dump (MHCII, CD11b, gdTCR, F4/80, CD19, NK1.1, CD11c)) plus one of the following Hashtag antibodies for each mouse: TotalSeq™-C0301, -C0302, -C0303, -C0305, -C0306, -C0307, C0308, -C0309, or -C0310. Cells from tumor or dLN were pooled by treatment, and viable CD8^+^ cells were sorted using a BD FACSAria™ Fusion flow cytometer (BD Biosciences, San Jose, CA). For TCR/scRNA-seq, the same downstream process described in the ‘*CD4 T cell scRNAseq analysis’* was followed. For scRNA/TCRseq, preparation of samples for loading to the 10X chip as well as reagents and instruments was performed as described in the CD4^+^ T cell library preparation section above.

#### Pre-processing of single-cell data

Sequencing files from Illumina assays were run through Cell Ranger v6.1.1 against a transcriptome derived from Ensembl v2.2.0 for the mouse genome GRCm38. For samples of CD4^+^ T cells, which were not multiplexed, matrix files from the filtered_feature_bc_matrix directory for RNA expression were converted into matrices and Seurat objects. UMI counts for Ig and TCR alleles were combined into a single value for the corresponding gene (e.g., allele counts *Trbv1* through *Trbv31* were combined into a single value for the *Trbv* gene). For the multiplexed samples of CD8^+^ T cells and CD45^+^ cells, matrix files were demultiplexed based on the highest count value among the 10 barcodes used for multiplexing, and the resulting sample-specific matrices were converted into Seurat objects, also combining Ig and TCR allele values into corresponding genes, as described previously. Data for each sample were processed using the scDblFinder package v1.16.0 to remove doublets. For samples of CD4^+^ and CD8^+^ T cells, TCR sequence data from the filtered_contig_annotations.csv files were processed using a custom script that identified clones across multiple tissues (spleen and tumor for CD4^+^ T cells; lymph node and tumor for CD8^+^ T cells), based on identical sets of alpha and beta sequences. For the CD45^+^ cells from tumor, the same procedure was performed on the TCR sequence data albeit for only a single tissue, as well as an analogous process of cross-tissue clone matching for the BCR sequences for the CD45^+^ cells from tumor and the B cells from the spleen. Subsequent analyses were performed as described below using Seurat package version 5.0.1 and the R statistical language version 4.3.1.

#### Processing of CD4^+^ T single cells

Matching samples from the spleens and tumors of the 6 mice, 3 in each experimental group, were combined into a single Seurat object using the FindIntegrationAnchors, IntegrateData, and ScaleData procedures with default parameters. This integration procedure from Seurat version 4 was used because our analysis predated the release of Seurat version 5, although Seurat 5 fully supports these procedures, and all scripts were re-run using Seurat 5. Cells were clustered using RunPCA and RunUMAP in 30 dimensions, and then FindNeighbors and FindClusters with a resolution of 0.5. Analysis of this combined dataset revealed that mouse S13 from the envRV group contributed to a tumor cluster not observed in any other tumor sample, and a spleen sample with a large proportion of dying cells, so that mouse was removed from the dataset. In addition, the UMAP reduction showed two large divisions with parallel sets of clusters that appeared to be explained primarily by expression of Ptprc. Therefore, another round of scaling and clustering was performed with the same parameters as before, but excluding mouse S13 and regressing out the integrated expression value of Ptprc using the vars.to.regress parameter in ScaleData.

Cluster identification: The Tfh cluster was defined based on expression of *Bcl6*, *Icos*, *Il21*, *Pdcd1*, *Tcf7*, and *Sostdc1,* while Th1 clusters were identified by *Tbx21*, *Cxcr6*, *Cxcr3*, *Id2*, *Ccl5*, *Runx3*, and *Nkg7* expression. One cluster termed Itg contained high expression of integrins, such as *Itga1* and *Itgb2*. The Treg cluster expressed *Foxp3*, *Il2ra,* inhibitory molecules *(Ctla4* and *Tigit)*, as well as the MHCII invariant chain *Cd74*. A cluster termed Cd74 also had high expression of this gene and *H2-Ab1*. Naive cells were characterized by the expression of expected markers, such as *Sell*, *Lef1*, *Tcf7*, and *Ccr7*, and by the high expression of ribosomal genes. A closely related cluster Ribo had high expression of ribosomal genes but not naive markers. Finally, we observed cell clusters with mitotic phenotypes, and other clusters with high levels of mitochondrial genes suggestive of cell death (see also Figure 3A-B).

#### Processing of CD8^+^ T single cells

For one mouse from the envRV group and one mouse from the 2W1S group, no lymph node sample was available. Otherwise, matching samples from lymph nodes and tumors were available for each of the 3 mice in each of the four experimental groups. After demultiplexing and removal of doublets, as described above, all available samples from lymph nodes and tumors of mice were combined into a single Seurat object, using the FindIntegrationAnchors, IntegrateData, and ScaleData procedures with default parameters. Data were dimensionally reduced using RunPCA and RunUMAP in 30 dimensions, and then clustered with FindNeighbors and FindClusters originally with a resolution of 2.0, to identify potential contaminating clusters. One outlier cluster was identified as containing primarily red blood cells, as indicated by high expression of hemoglobin A (gene Hba-a1) and biphosphoglycerate mutase (Bpgm), and removed in a second round of normalization with SCTransform; dimensional reduction with RunPCA and RunMAP, and clustering with FindNeighbors and FindClusters, this time with a resolution of 0.3.

#### Separation of intratumoral CD45^+^ single cells

All 9 tumor samples and matching 9 spleen samples were combined into a single Seurat object, with the demultiplexing and double removal procedures as discussed above, and then the FindIntegrationAnchors, IntegrateData, and ScaleData procedures with default parameters. Cells were clustered using RunPCA and RunUMAP in 30 dimensions, and then FindNeighbors and FindClusters with a resolution of 2.0, a value chosen to help identify relatively small clusters that were homogeneous for a particular T cell subtype, B cell subtype, or other cell subtype. For each of the resulting 47 clusters, we performed another round of scaling and clustering at a resolution of 0.2, to determine if the cluster was indeed homogeneous, or could be separated into T cells, B cells, or other cells, based on the presence of BCR or TCR clonotypes and the markers Cd3d, Cd79a, Cd14, and Ccl22. The co-occurrence across cell types was frequently observed within each cluster. This process yielded a separation of cells into T, B, and other categories, which were then analyzed separately as described below.

#### Analysis of T single cells from the CD45^+^ cell dataset

We renormalized and rescaled the subset of T cells using NormalizeData, FindVariableFeatures, ScaleData, and SCTransform, using default parameters. We clustered using RunPCA, RunUMAP, and FindNeighbors, using default parameters, and FindClusters with a high resolution of 2.0, to separate CD8^+^ and CD4^+^ T cells. We used the markers CD8a and CD4 to assign clusters into CD8^+^ and CD4^+^ T cell categories, respectively. Three clusters needed further subclustering at a resolution of 0.2 to separate CD8^+^ and CD4^+^ T cells. Because of the relatively few T cells obtained in this dataset, we did not try to characterize them by clustering, but relied instead upon the clustering obtained from our large dataset of CD4^+^ T cells. We used the functions FindTransferAnchors using the LogNormalize method and TransferData using the protocol published by Seurat, to project the CD4^+^ T cells from the CD45^+^ cell dataset onto the clustering from the CD4^+^ T cell dataset. Analysis revealed that cells projecting onto the Th1.1 cluster showed a spatial separation in the UMAP plot based on their experimental group, suggesting that they had distinct phenotypes. We therefore applied the FindSubCluster procedure with a resolution of 0.08, chosen to produce two subclusters, named as Th1.1a and Th1.1b.

#### Analysis of B single cells from the CD45^+^ cell dataset

We renormalized and rescaled the subset of B cells using SCTransform with default parameters. We performed dimensionality reduction using RunPCA, RunUMAP in 20 dimensions because the default value of 30 subsequently resulted in several clusters that were split spatially in the UMAP projection. We performed clustering using FindNeighbors, using default parameters, and FindClusters with a resolution of 0.3.

#### Heatmaps of single-cell clusters

Heatmaps were computed on all single-cell clusterings to help with their characterization. For the CD4^+^ and CD8^+^ T cell datasets, in which integration was the last processing step, we used expression from the scaled data from the integration step. For the analyses of the CD45 datasets, in which subsetting, renormalization, and rescaling were the final processing steps, expression was taken from the scaled data of the RNA count matrix. Expression values were converted into Z-scores for each gene across samples, and centroids were computed for each cluster by taking the trimmed mean of each Z-score across the cluster, removing the top and bottom 10 percent of values. Biomarkers for each cluster were identified among genes with the highest coefficients of variation across centroid values. Heatmap figures were generated using the superheat package.

### Adoptive T cell transfer experiments

Three successive cohorts of E0771 tumor-bearing mice were vaccinated i.v. with envRV RNA-LPX on days 7, 14 and 21 then spleen and tumor tissues were collected on day 26. 95,000-135,000 envRV^+^ CD4^+^ T cells sorted from tumor and spleen and 82,000-85,000 CD8^+^ T cells sorted from tumor were transferred intratumorally in 50uL PBS into WT or Batf3^-/-^ recipient E0771 tumor-bearing mice on days 1, 4 and 7 after tumor implantation. Single vaccine or combination groups received the same number of envRV^+^ CD4^+^ T cells and CD8^+^ T cells. PBS was injected intratumorally in the control group.

### ELISPOT

5x10^5^ CD4-depleted or CD8-depleted splenocytes (using the manufacturer’s procedure with CD4 L3T4 microbeads or CD8 Ly-2 microbeads (Miltenyi, Cat#130-117-043, Cat#130-117-044) and LS columns (Miltenyi, Cat#130-042-401) were cultured overnight at 37°C in RPMI 1640 containing 10% FBS (1% penicillin/streptomycin, 1% Hepes, 1% GlutaMAX, and 1% sodium pyruvate) in a 96-well mouse IFN-γ ELISpot plate (Biotechne, Cat#EL485). CD4 and CD8-depleted splenocytes (containing DCs) were used to determine the CD8^+^ and CD4^+^ T cell response respectively in an ELISpot assay, and only samples with <2% of remaining depleted cells were considered in the analysis. For stimulation, each long peptide (envRV and 2W1S) was added into the culture at 10 μg/ml and short peptide (envRV-E) at 1 μg/ml. Restimulation with CD3/CD28 Dynabeads (Gibco, Cat#11452D) or vehicle (DMSO, Sigma Cat#D2650) were used as controls. For analysis, IFN-γ spots were counted using the manufacturer’s procedure (R&D Systems mouse IFN-gamma kit) and an automatic ELISpot reader (AID). All samples were tested in 3 biological replicates. Peptide synthesis was performed by Genscript (75% purity).

### Detection of envRV-specific antibodies by ELISA

EnvRV protein (produced in-house) or envRV 29mer (Genscript) were coated in flat-bottom 384-well plates (Nunc Maxisorp 384-well plates (Cat #464718)) at 1 μg/ml in PBS at 4°C overnight. The day after, plates were washed four times with PBS 0.05% Tween-20 before incubation in PBS (no Ca no Mg) 0.5% BSA for 2 hours at room temperature for the blocking step. Plates were then washed four times with PBS 0.05% Tween-20 before incubation for 1 hour and 20 minutes at room temperature with serum from mice immunized with envRV RNA-LPX or 2W1S RNA-LPX vaccines diluted in PBS 0.05% Tween-20 0.5% BSA. Plates were then washed five times with PBS 0.05% Tween-20 before incubation for 1 hour at room temperature with Goat anti-mouse Immunoglobulin conjugated with horseradish peroxidase (SouthernBiotech) (anti-mouse total IgG(H+L)-HRP (Cat #1036-05), anti-mouse IgA-HRP (Cat #1040-05), anti-mouse IgM-HRP (Cat #1021-05), anti-mouse IgG1-HRP (Cat #1071-05), anti-mouse IgG2c-HRP (Cat #1078-05)) diluted 1/2000 in PBS 0.05% Tween-20 0.5% BSA. Plates were then washed five times with PBS 0.05% Tween-20 before incubation for 15 minutes at room temperature with TMB solution (Thermo Scientific Cat#34028). An equivalent volume of H_2_SO_4_ 0.1M solution was added to stop the reaction. Plate reading was done on (Agilent Biotek Synergy H1) at 450nm.

### Anti-envRV antibody generation

C57BL/6 mice were vaccinated with 50 μg envRV RNA-LPX by i.v. injection on days 7, 14, 21 after tumor implantation followed by peptide vaccine 100 μg/ml by s.c injection on days 28 and 35. Three days after the last dose, B cells from the lymph nodes and spleens of immunized mice were enriched as previously described^61^. Enriched cells were stained with biotinylated envRV at 5 ug/ml followed by streptavidin APC (Biolegend 405207, 1: 100). EnvRV-specific B cells were sorted and cultured as previously described^61^. The single B cell culture supernatants were screened for envRV and anti-mouse IgG (Rockland) binding clones, the positive clones were screened by flow cytometry against E0771 cells. B cells producing envRV-reactive IgGs were picked to generate recombinant monoclonal antibodies as previously described^61^. In the serum of E0771 tumor-bearing animals vaccinated with envRV RNA-LPX, IgG1 and IgG2c were the primary antibody subtypes detected. IgG2c/IgG2a are the isotypes exhibiting the most potent effector functions (ADCC, ADCP, CDC), in comparison to IgG1. In C57BL/6 mice, IgG2c is expressed instead of IgG2a. However, since IgG2a is a more common isotype and facilitates quick monoclonal antibody production using existing hybridoma/cell lines, along with the availability of isotype controls, we opted to generate the specific monoclonal antibody using the IgG2a isotype.

### E0771 *in vitro* binding assay

E0771 tumor cells were harvested using 25mM EDTA solution and 5x10^5^ cells were seeded in round bottom 96-well plates and incubated for 20 minutes at 4°C with 5μg/ml of mouse FcBlock (BD Cat#553141) and 1:100 of LIVE/DEAD Fixable Near-IR (ThermoFisher Cat#L10119) in 50μl of HBSS (Cat#14-025-092). After washing with HBSS, cells were incubated with ten-fold serial dilutions of serum from C57BL/6 E0771 tumor-bearing mice previously vaccinated with 50 μg envRV or 2W1S RNA-LPX on days 7, 14 and 21 after tumor implantation. When indicated, E0771 tumor cells were incubated with serial dilutions of in-house-generated monoclonal anti-envRV antibody. After washing with HBSS twice, cells were then incubated with PE-conjugated Goat anti-mouse IgG (BioLegend Cat#405307) 1/200 diluted in HBSS for 20 minutes at 4°C. Cells were washed twice with HBSS before acquisition on a Symphony flow cytometer (BD Biosciences).

### *In vivo* anti-envRV antibody treatment experiment

Anti-envRV monoclonal antibody (ɑ-envRV mAb) or in-house mouse ɑ-GP120 IgG2a isotype control were administered i.v. in C57BL/6 E0771 tumor-bearing mice on days -6, -2, 2, 6, 9, 12, 14, 16, 20, 22, 26 and 29 at 400 μg/100μl/mouse. Tumors were measured twice weekly using digital calipers, and tumor volumes calculated using the modified ellipsoid formula, 1/2 x (length x width^2^). Mice were euthanized if tumors ulcerated or volumes exceeded 1500 mm^3^.

### Cytokines/chemokines serum analysis

RO bleeding was performed on mice vaccinated i.v with the RNA-LPX vaccine 6 hours before. Blood was collected in Serum-Gel Polypropylene tubes (BD Microtainer SST #365967) and spin at 6,000 rpm for 5 min at room temperature. The clear fraction (serum) was collected, frozen at -80C and thawed just before the cytokine assay. Serum samples were analyzed for concentrations of IL-12p40, TNFa, CCL5, CXCL9 and CXCL10 using Milliplex MAP reagents (MilliporeSigma, #SPRMCYPMX32-BK) according to the manufacturer’s recommended protocol. Dilutions of reconstituted standards were made at 2.73-fold to increase the number of data points from six to nine while maintaining the original dynamic range. Fluorescence intensity was measured by xPONENT software v 4.2 on FlexMap 3D instruments (Luminex Corp, Austin, TX). Bio-Plex Manager v 6.2 (Bio-Rad Laboratories, Hercules, CA) was used to construct standard curves for each analyte on each plate using the median FI of at least 20 beads done in duplicate. A 4 or 5- point regression analysis was used to calculate best fit and cytokine levels.

### Tumor histology

Immunohistochemistry for F4/80 (CI:A3-1, Serotec, cat no. MCA497G) and NK1.1 (E6Y9G, Cell Signaling Technologies, cat no. 29197S) was performed on 4 μm-thick paraffin-embedded tissue sections mounted on Superfrost Plus glass slides. Tissue sections were deparaffinized and rehydrated using deionized water. F4/80 (10 μg/mL) was incubated on sections for 60 min at room temperature following Target pH 6.0 antigen retrieval. Subsequently, endogenous peroxidase was quenched by incubating sections in 3% H_2_O_2_ for 4 min at room temperature. F4/80 was detected using anti-rat (7.5 μg/mL) secondary antibody with the ABC Peroxidase Elite Detection Kit (Vector Laboratories, Burlingame CA; Cat #PK-6100). For NK1.1, Ventana CC1 standard antigen retrieval was performed on sections according to the manufacturer’s instructions. NK1.1 (0.125 μg/mL) was incubated on sections for 60 min at 37C followed by Ventana rabbit OmniMap detection. Sections were counterstained with Mayer’s hematoxylin, dehydrated, mounted with permanent mounting medium and cover-slipped.

### Statistical analysis

Unless otherwise stated (e.g. for scRNAseq analysis), GraphPad Prism 10 was used for data analysis and representation. The ANOVA statistical test with multiple comparisons tests were used unless otherwise stated in figure legends. All tests were two-sided with a significance threshold of 0.05. Summary line graphs, bar charts, and associated data points represent means of data; error bars indicate SEM.

## Acknowledgements

We thank the members of the Genentech Cancer Immunology department for discussions and experimental assistance; the Genentech vivarium facility staff for animal care and maintenance; colleagues of the Genentech sequencing, Protein Production Group, and flow cytometry core laboratories; Nathan West and Christine Moussion for relevant feedback on the manuscript; Jieming Chen and Daniel Oreper for bioinformatics analysis support; Cecile de la Cruz for assistance with outsourced *in vivo* studies; the Genentech Postdoc Program for scientific training and support. Illustrations were created in BioRender. This study was funded by Genentech.

## Author contributions

Conceptualization, A-H.C., I.M., and J.L.L.; design and methodology, A-H.C, C-C.B., R.B., J.L., T.W., S.D., M.S., D.S., B.B., J.Z., E.F., A.E., J.C.; data acquisition, A-H.C, C-C.B., R.B., J.L., J.G., C.C., A.N., M.Y., A.G., Y.C., V.J., H.M., Y.S., M.P., T.J., Y.W., L.L., E.D., A.A., A.A., Y.O., J.C., S.L., J.P., J.H., S.W., J.O., S.T., and J.L.; data analysis and interpretation, A-H.C, C-C.B., R.B., T.W., and J.L.; writing – original draft, A-H.C, C-C.B., J.L., and T.W.; writing – review and editing, L.D., I.M., and R.B.; writing - final review, all authors.

## Supplemental information

**Figure S1:**
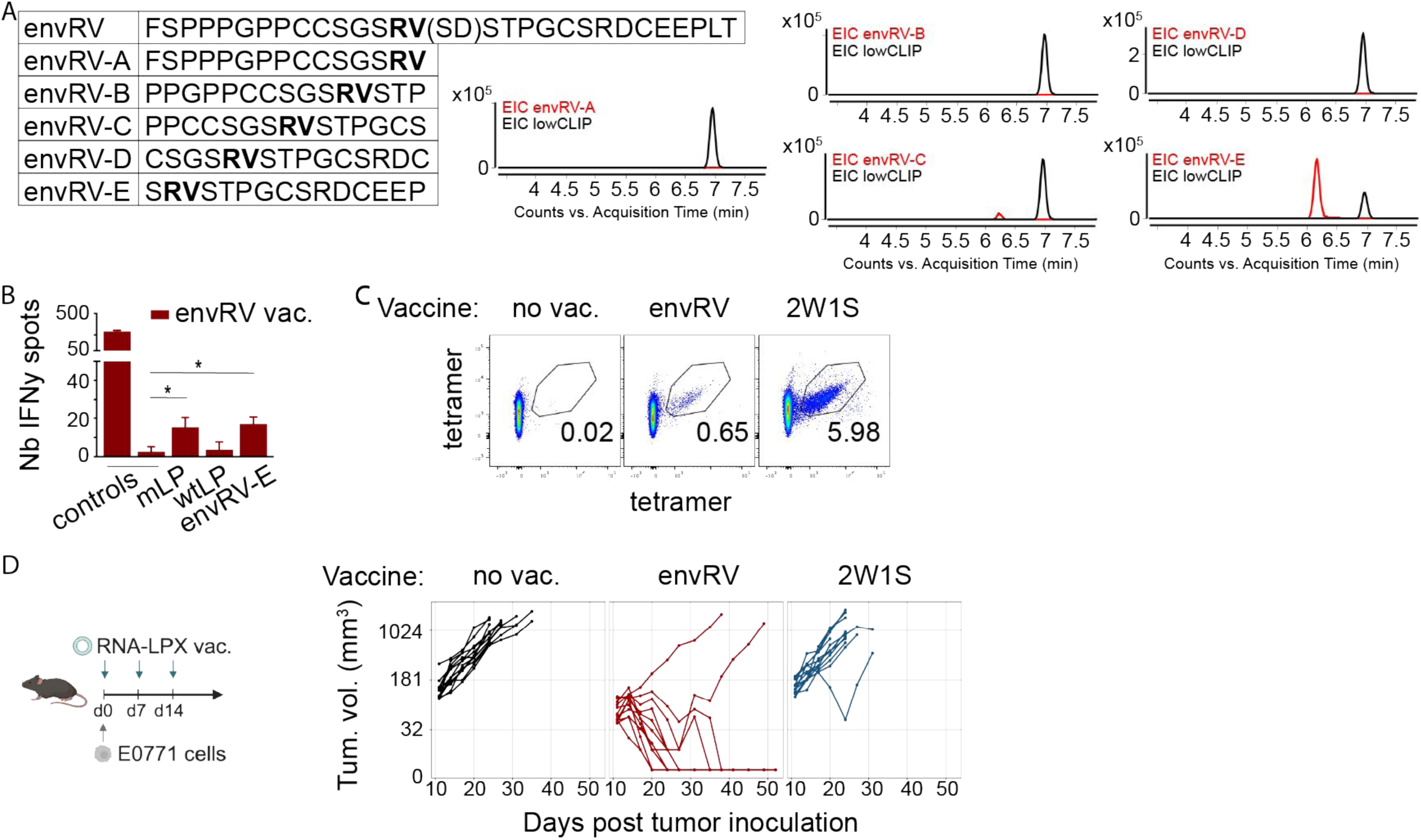
Detection of envRV:I-A^b^-specific CD4^+^ T cells and tumor rejection. A: Peptide library of overlapping 15 mers for the envRV mutated sequence tested for MHC-II binding with the 2D-LCMS assay. The double point mutation is represented in bold and the WT amino-acid (AA) shown in parenthesis. Graphs with curve for each peptide of interest (red curve) and lowCLIP peptide control (black curve) are shown. B: Mouse IFN𝛾 ELISpot of splenocytes depleted of CD8^+^ T cells (CD4 response) from vaccinated mice with envRV long peptide (29 AA). 5x10^5^ splenocytes were restimulated overnight with either the envRV long peptide (29AA, mLP), the WT counterpart long peptide (29 AA, wtLP) or the neoepitope envRV-E peptide (15 AA) identified in A. CD3/CD28 beads were used for positive control, and absence of peptide (dmso vehicle) for negative control. Data of the number of IFN𝛾 spots are represented as mean +/- SD. *p<0.05. C: EnvRV-E and 2W1S -specific tetramers were used to detect envRV+ and 2W1S+ CD4^+^ T cells respectively in spleens of vaccinated mice with envRV or 2W1S RNA-LPX. Non-vaccinated mice were used as negative control. Representative FACS plots are shown for each condition. D: E0771 tumor-bearing mice were either vaccinated with envRV (neoantigen) or 2W1S (irrelevant tumor antigen) RNA-LPX starting on the day of tumor inoculation, for a total of 3 vaccines administered 7 days apart. Non-vaccinated animal were used as control. Volume of tumors were measured for 52 days post tumor inoculation.

**Figure S2:**
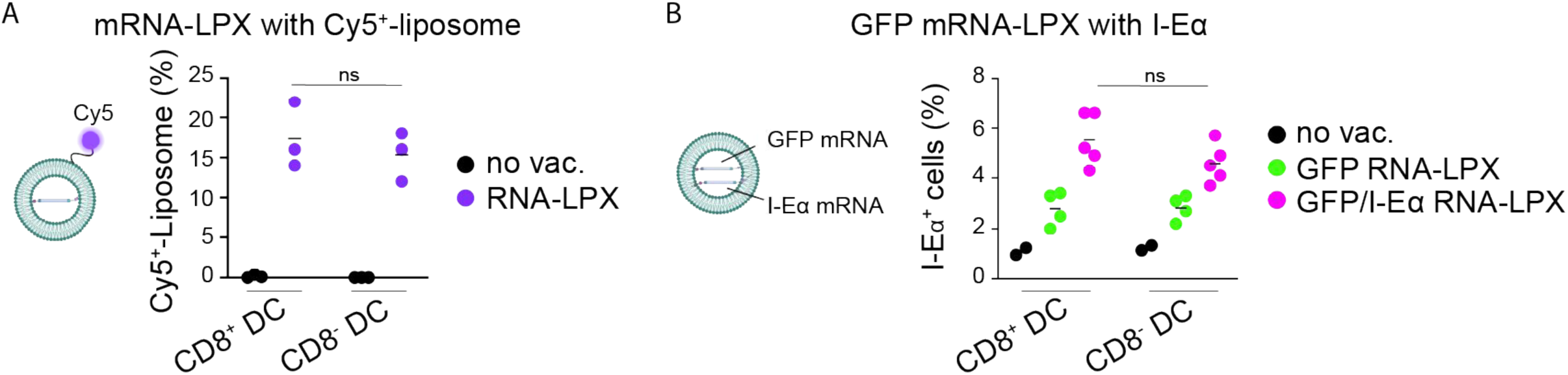
cDC1 and cDC2 uptake and present mRNA-LPX encoded antigens. A: Mean proportion of Cy5^+^-liposome uptaken by CD8^+^ and CD8^-^ DCs from spleen of WT mice following a single vaccination and measured by flow cytometry. Spleens were harvested one hour post vaccination or in the absence of RNA-LPX vaccine. ns: non significant. B: Mean proportion of YAe (I-Eɑ_52-68_:I-Ab)^+^ presented by CD8^+^ and CD8^-^ DCs from spleen of WT mice following a single vaccination and measured by flow cytometry. Spleens were harvested seventeen hours post vaccination or in the absence of RNA-LPX vaccine. Mice received GFP or a combination of GFP and I-Eɑ_52-68_ RNA-LPX vaccine. ns: non significant.

**Figure S3:**
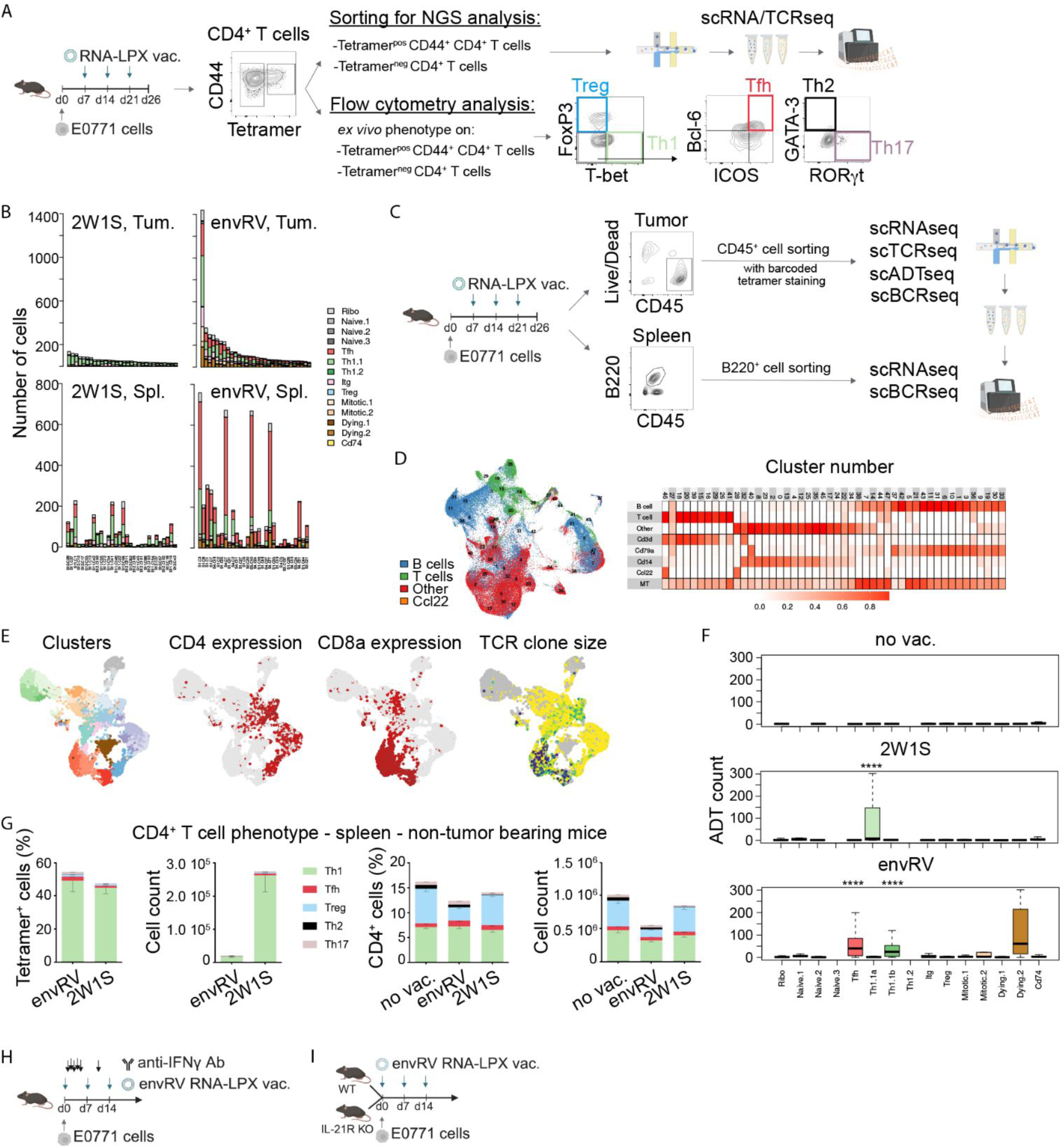
Tumor influences the phenotype of vaccine-induced CD4 T cells A: Study design and gating strategy for cell sorting for NGS analysis, and *ex vivo* flow cytometry analysis. Spleens and tumors of mice vaccinated with envRV or 2W1S RNA-LPX were harvested 5 days post last vaccination, and single cell suspensions were stained to define lymphoid lineages and envRV or 2W1S IA^b^ tetramer-positive CD4^+^ T cells. For the NGS experiment, tetramer^pos^CD44^+^ CD4^+^ T cells and tetramer^neg^ CD4^+^ T cells were sorted separately from spleen and tumor. For each organ, sorted cells (tetramer^pos^ and tetramer^neg^) were then mixed at equal ratio and submitted to scRNA/TCRseq analysis (n=3 spleens and n=3 tumors for each treatment group, for a total of n=12 samples). For *ex vivo* phenotypic analysis by flow cytometry in subsequent studies, cells from spleen and tumor were stained with TFs to define canonical CD4^+^ T cell phenotypes (T-bet^+^ FoxP3^-^ cells for Th1, FoxP3^+^ T-bet^-^ cells for Treg, and from the FoxP3^-^ T-bet^-^ population, Bcl-6^+^ICOS^+^ cells for Tfh, GATA-3^+^ cells for Th2, and ROR𝛾t^+^ cells for Th17. B: Barplot graphs representing the number of cells and phenotype of matching clones in tumors (top panels) and spleens (lower panels) from 2W1S (left panels) and envRV (right panels) RNA-LPX vaccinated animals. Phenotypes are colored according to the UMAP in Figure 3A. C: Study design and gating strategy for cell sorting of the experiment shown in Figures 3F, 4C-I, S4B-C. Tumors and spleens were isolated from mice vaccinated with envRV or 2W1S RNA-LPX. Non-vaccinated tumor-bearing animals were used as control. Tumor cells were stained with barcoded envRV:IAb tetramer (non-vaccinated and envRV RNA-LPX groups) or 2W1S:IAb tetramer (2W1S vaccine group), then CD45^+^ cells were sorted and submitted to scRNA/TCR/ADTseq; for further analysis of the tetramer^+^ CD4^+^ T cells (used in Figure 3F), and scRNA/BCRseq; for further analysis of the intratumoral B cell compartment (used in Figures 4 and S4). In parallel, B220^+^ cells were sorted from spleens of the 3 treatment groups and submitted to scRNA/BCRseq for further analysis of splenic B cells (used in Figures 4 and S4). D: UMAP of the whole dataset combining CD45^+^ cells from tumors and B220^+^ cells from spleens described in C. Heatmap depicting cluster numbers and their categorization as B, T or other cells based on CD3d, CD79a, CD14 and CCl22 expression. E: UMAPs of the T cell subsets identified in D, and further differentiating CD4^+^ and CD8^+^ T cell clusters as well as TCR clone sizes. Computationally isolated CD4^+^ T cells were then used for the CD4 transfer analysis in Figures 3F and S3F. F: Barplots depicting ADT counts for each phenotype of the CD4^+^ T-cell transfer analysis from the CD45^+^ cell dataset to the CD4 single-cell dataset described in Figure 3 A-E. Data are represented as boxplots with median for each treatment group. ****p<0.0001. G: Phenotypic analysis of CD4^+^ tetramer^pos^ cells (left panel) and tetramer^neg^ cells (right panel) in spleen from envRV or 2W1S vaccinated non-tumor bearing animals. No vaccination was used as control. Flow cytometry data are shown as mean +/- SEM. H: Efficacy study design with IFN𝛾 depletion. E0771 tumor-bearing mice were vaccinated with envRV RNA-LPX starting at day 0 for a total of 3 vaccines given 7 days apart. Isotype control or anti-IFN-γ antibodies were administered at 1 mg IP then 0.5 mg IP on days 1,2,3, 4, 5, 11. See Figure 3H. I: Efficacy study with WT or IL21-R KO mice bearing E0771 tumor cells and vaccinated with envRV RNA-LPX starting on day 0 and every 7 days for a total of 3 vaccinations. See Figure 3I.

**Figure S4:**
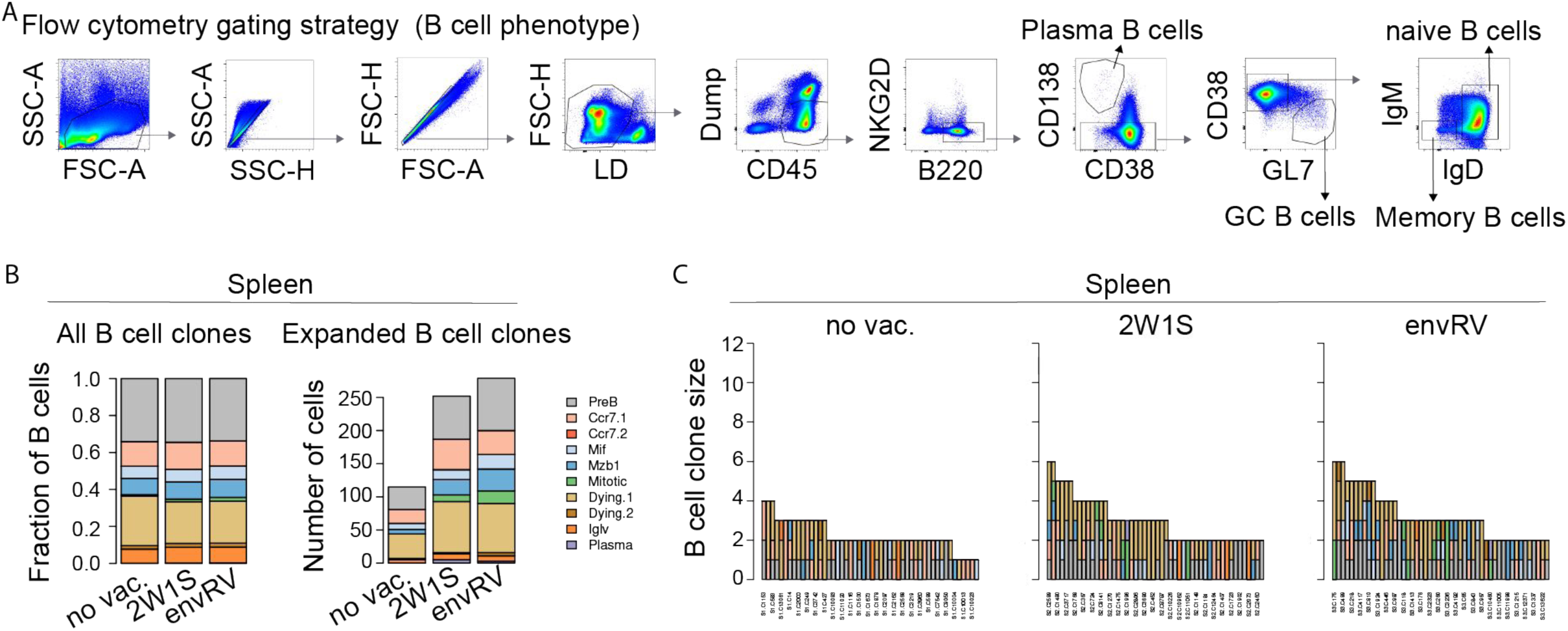
B-cell phenotype in spleen. A: Flow cytometry gating strategy for identifying B cell subsets in spleen and tumor for Figure 4B: Plasma B cells (Live^+^B220^+^CD138^+^CD38^-^), Germinal Center (GC) B cells (Live^+^B220^+^CD138^-^CD38^-^GL7^+^), naive B cells (Live^+^B220^+^CD138^-^CD38^+^GL7^-^IgD^+^IgM^+^), and memory B cells (Live^+^B220^+^CD138^-^CD38^+^GL7^-^IgD^-^IgM^-^). Live LD: Live dead staining; dump channel: CD3, CD11b, F4/80, CD11c. B: Stacked bar graphs of B cell cluster composition (as shown in Figure 4D) of all B cells (left) and expanded clones (right) in spleen from non-vaccinated (no vac.), control (2W1S) and envRV RNA-LPX vaccinated mice. C: Bar graph showing the primary B cell clonotypes in spleen from non-vaccinated (no vac.), control (2W1S) or envRV RNA-LPX vaccinated mice, colored as in Figure 4D. Legend is the same as in B.

**Figure S5:**
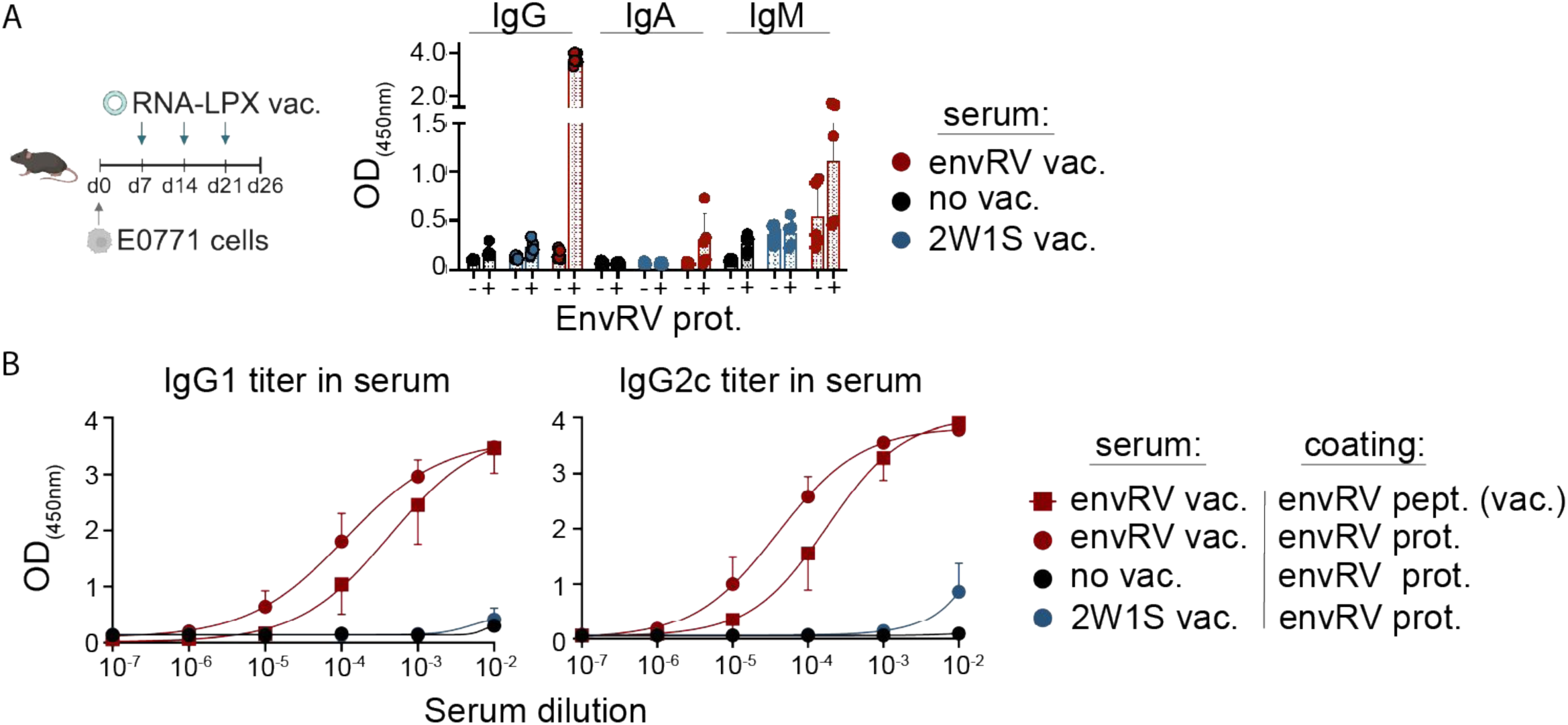
Isotype of envRV-specific antibodies in serum. A-B: E0771 tumor-bearing mice were vaccinated with envRV (neoantigen) or 2W1S (control) RNA-LPX starting 7 days post tumor inoculation for a total of 3 vaccines given 7 days apart. No vaccination was used as control. Serum was collected 5 days post last vaccination. (A) Recognition of the envRV full protein was determined by ELISA for the IgG, IgA and IgM isotypes. Data are shown as (OD_450nm_) values in presence or absence of the coated envRV protein. (B) IgG1 and IgG2c titers were determined by ELISA in serum from envRV-vaccinated mice recognizing the mutated full envRV protein (envRV prot.; red circle curve) or the mutated envRV peptide encoded in RNA-LPX vaccine (envRV pept. (vac.); red square curve). Serum from non vaccinated (black curve) or 2W1S RNA-LPX vaccinated animals (blue curve) were used as control. Data are shown as (OD_450nm_) values for dilution of serum from 10^-2^ to 10^-7^.

**Figure S6:**
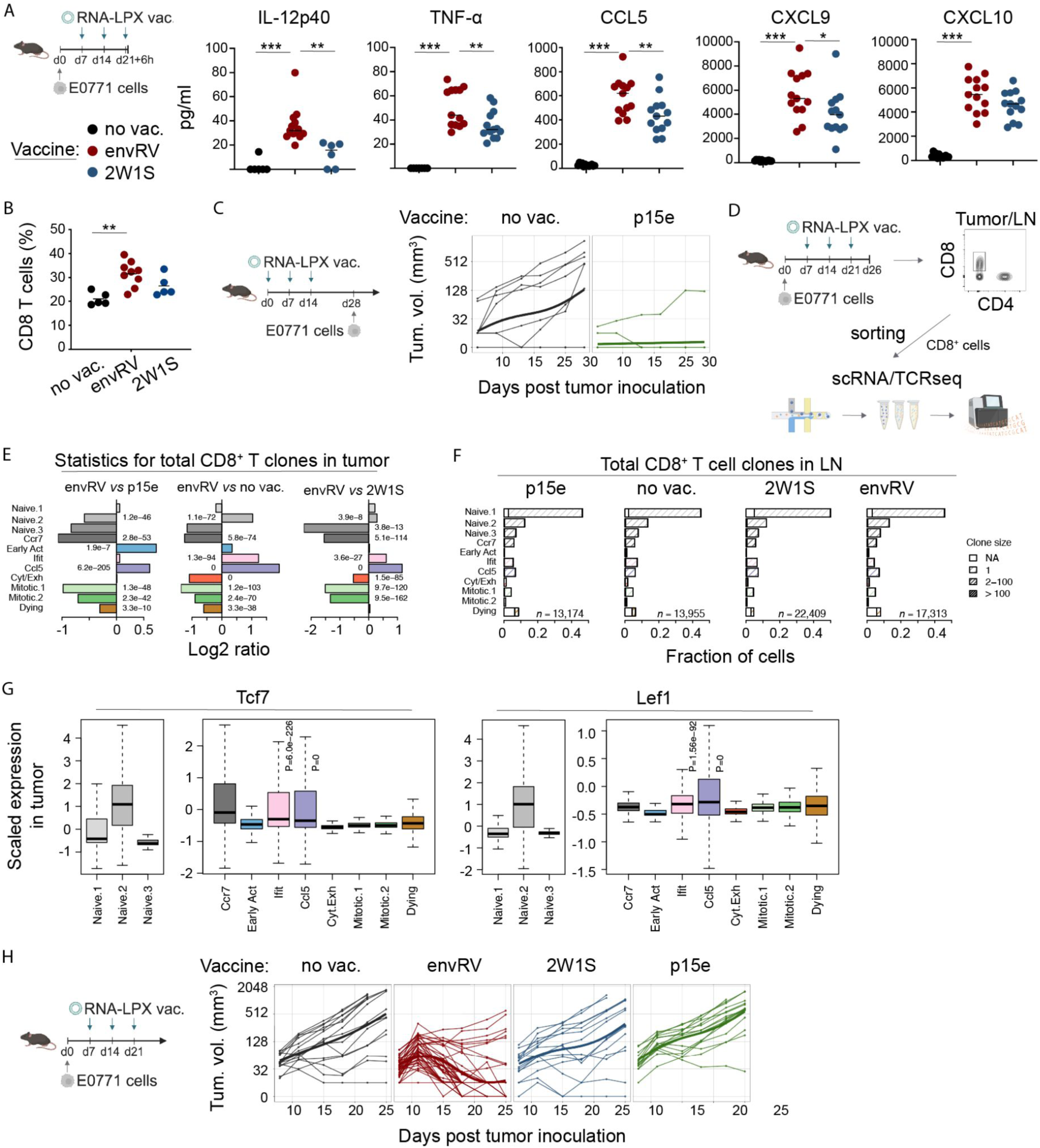
EnvRV RNA-LPX vaccine enhances CD8^+^ T cell infiltration in tumor and dLNs leading to tumor rejection, in contrast to the MHCI p15e vaccine. A: Serum levels of cytokines and chemokines 6 hours after the third vaccine dose of envRV or 2W1S RNA-LPX vaccine administered in tumor-bearing mice. Non vaccinated tumor-bearing mice were used as control (no vac.). Data shows three independent pooled experiments. Data are shown as mean.*p<0.05, **p<0.01, ***p<0.001. B: Proportion of CD8^+^ T cells among CD45^+^ cells in tumor-draining LNs analyzed by ex vivo staining on d26 post tumor inoculation. Non-vaccinated tumor-bearing mice were used as control. Representative data of two independent experiments. Data are shown as mean. **p<0.01. C: Prophylactic efficacy study with the MHCI p15e vaccine. Tumor growth curves of mice vaccinated or not with MHCI-restricted (p15e) RNA-LPX starting 28 days prior to tumor inoculation are shown. Mice were challenged 14 days post last vaccine with E0771 tumor cells. A total of 3 vaccines, 7 days apart, were administered. Non-vaccinated animals (no vac.) were used as control. D: Study design for scRNA/TCRseq analysis. E0771 tumor-bearing mice received or not three doses of envRV, 2W1S or p15e RNA-LPX vaccine. Five days post last vaccine tumor dLNs and tumors were harvested and CD8^+^ T cells were sorted for scRNA/TCR sequencing. E: Barplots showing the fraction of CD8^+^ T cells from LNs for each cluster identified from the UMAP in Figure 6C. In each stacked bar, open bar denotes cells for which TCR were unidentified, light stripes for singletons, medium stripes for clones between 2-100 cells, and high density stripes for clones greater than 100 cells. Stripe color represents the primary cell phenotype of each clone as shown in the UMAP. F: Barplots showing the log2 ratio of the total fraction of CD8^+^ T cells for each phenotypic cluster in tumors. Comparisons between envRV and each of the 3 other treated groups with significant statistical values are shown. G: Boxplot graphs showing the scaled expression in tumor of the Tcf7 (left) and Lef1 (right) genes for each clusters identified in the UMAP in Figure 6C. Statistical significant values from each clusters versus all other clusters (but naive clusters) are indicated on the graphs. H: Study design of the efficacy study with the p15e vaccine given in a therapeutic setting. E0771 tumor-bearing mice were vaccinated either with MHCII-restricted (envRV or control 2W1S) or MHCI-restricted (p15e) RNA-LPX starting 7 days post tumor inoculation. A total of 3 vaccines, 7 days apart, were administered. Non-vaccinated animals (no vac.) were used as control.

**Figure S7:**
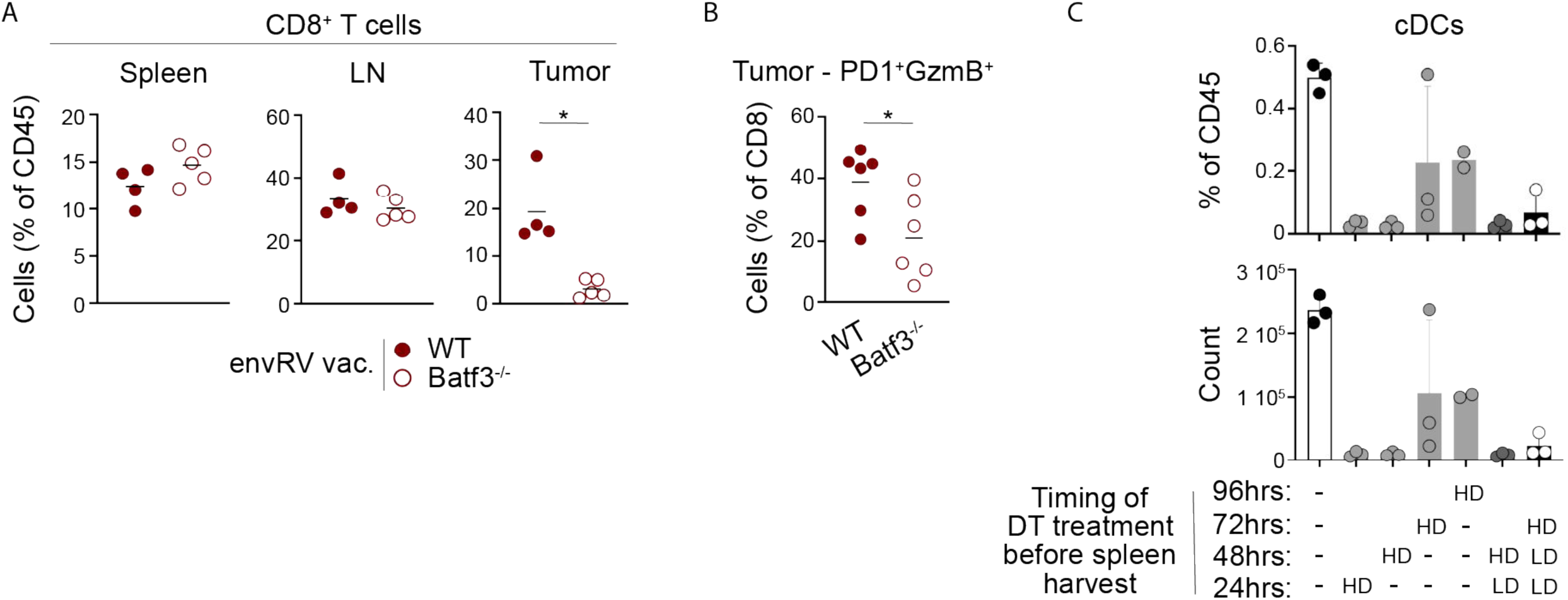
cDC1s are critical for CD8^+^ TIL accumulation and cytotoxicity following envRV RNA-LPX vaccination. A: Frequencies of CD8^+^ T cells among CD45^+^ cells in spleen, tdLN and tumor from WT and Batf3^-/-^ mice 5 days following the third dose of envRV RNA-LPX vaccine. Mean data are shown from one representative experiment out of 3 independent ones. Normality and Mann-Whitney or t-tests were used. *p<0.05. B: Frequencies of PD-1^+^GzmB^+^ CD8^+^ TILs from WT and Batf3^-/-^ mice 5 days following the third dose of envRV RNA-LPX vaccine. Mean data are shown from one experiment. Normality and Mann-Whitney or t-tests were used. *p<0.05. C: Proportion and count of cDCs identified by flow cytometry from splenocytes of mice treated or not with one single high dose (HD, 500ng) of dyphteria toxine (DT) at 24, 48, 72 or 96 hours prior to takedown (TD); or treated with one single HD of DT at 48h prior TD, followed by one single low dose (LD, 100ng) of DT at 24h prior TD; or treated with one single HD of DT at 72h prior TD, followed by two LD of DT at 48 and 24h prior TD. Data are shown as mean +/- SD.

